# Two promoters integrate multiple enhancer inputs to drive wild-type *knirps* expression in the *D. melanogaster* embryo

**DOI:** 10.1101/2021.03.23.436657

**Authors:** Lily Li, Rachel Waymack, Mario Elabd, Zeba Wunderlich

## Abstract

Proper development depends on precise spatiotemporal gene expression patterns. Most genes are regulated by multiple enhancers and often by multiple core promoters that generate similar transcripts. We hypothesize that these multiple promoters may be required either because enhancers prefer a specific promoter or because multiple promoters serve as a redundancy mechanism. To test these hypotheses, we studied the expression of the *knirps* locus in the early *Drosophila melanogaster* embryo, which is mediated by multiple enhancers and core promoters. We found that one of these promoters resembles a typical “sharp” developmental promoter, while the other resembles a “broad” promoter usually associated with housekeeping genes. Using synthetic reporter constructs, we found that some, but not all, enhancers in the locus show a preference for one promoter. By analyzing the dynamics of these reporters, we identified specific burst properties during the transcription process, namely burst size and frequency, that are most strongly tuned by the specific combination of promoter and enhancer. Using locus-sized reporters, we discovered that even enhancers that show no promoter preference in a synthetic setting have a preference in the locus context. Our results suggest that the presence of multiple promoters in a locus is both due to enhancer preference and a need for redundancy and that “broad” promoters with dispersed transcription start sites are common among developmental genes. Our results also imply that it can be difficult to extrapolate expression measurements from synthetic reporters to the locus context, where many variables shape a gene’s overall expression pattern.

## Introduction

Diverse processes in biology, from early development to the maintenance of homeostasis, rely on the regulation of gene expression. Enhancers and promoters are the primary regions of the genome that encode these gene regulatory programs. Both enhancers and promoters are characterized by clusters of sequence motifs that act as platforms for protein binding, allowing for the integration of a spectrum of signals in the cellular environment. The majority of studies that dissect enhancer or promoter function typically investigate each in isolation, which assumes that their function is largely modular. In practice, this means that we assume an enhancer drives generally the same pattern, regardless of promoter, and that promoter strength is independent of the interacting enhancer. However, there is evidence that there can be significant “interaction terms” between promoters and enhancers, with enhancer pattern shaped by promoter sequence, and promoter strength influenced by an enhancer (Gehrig et al., 2009; Hoppe et al., 2020; Qin et al., 2010).

Therefore, a key question is precisely how the sequences of an enhancer and a promoter combine to dictate overall expression output. Adding to the complexity of this question, developmental genes often have multiple enhancers, and many metazoan genes have alternative promoters (Brown et al., 2014; Landry, Mager, & Wilhelm, 2003; Schibler & Sierra, 1987; Schröder, Tautz, Seifert, & Jäckle, 1988). In a locus, multiple enhancers exist either because they drive distinct expression patterns or, in the case of seemingly redundant shadow enhancers, because they buffer noise in the system (Kvon, Waymack, Elabd, & Wunderlich, 2021). Though RAMPAGE data shows that >40% of developmentally expressed genes have more than one promoter (P. Batut, Dobin, Plessy, Carninci, & Gingeras, 2013), the role of multiple promoters has been relatively less explored. In some cases, alternative promoters drive distinct transcripts, but *hunchback* is a notable example of a gene with two highly conserved promoters that produce identical transcripts (Ling, Umezawa, Scott, & Small, 2019; Schröder et al., 1988).

This suggests there may be additional explanations for the prevalence of multiple promoters. One possibility is molecular compatibility—promoters can preferentially engage with different enhancers depending on the motif composition and proteins recruited to each (van Arensbergen, van Steensel, & Bussemaker, 2014; Wang, Hou, Quedenau, & Chen, 2016). For example, enhancers bound by either the transcription factors (TFs) Caudal or Dorsal tend to interact with Downstream Promoter Element (DPE)-containing promoters (Juven-Gershon, Hsu, & Kadonaga, 2008; Zehavi, Kuznetsov, Ovadia-Shochat, & Juven-Gershon, 2014) and Bicoid-dependent *hunchback* transcription seems to depend on the presence of a TATA box and Zelda site at one promoter (Ling et al., 2019). Another possibility is that having multiple promoters provides redundancy needed for robust gene expression, much like shadow enhancers.

To distinguish between these hypotheses, an ideal model is a gene with (1) multiple promoters that contain different promoter motifs and drive similar transcripts and with (2) multiple enhancers bound by different TFs. The *Drosophila* developmental gene *knirps* (*kni*) fits these criteria. It is a key developmental TF that acts in concert with other gap genes to direct anterior-posterior axis patterning of the early embryo. *Kni* has two core promoters that drive nearly identical transcripts (only differing by five amino acids at the N-terminus) and that are both used during the blastoderm stage (Figure 1A – C). Here, we define the core promoter as the region encompassing the transcription start site (TSS) and the 40bp upstream and downstream of the TSS (Vo Ngoc, Wang, Kassavetis, & Kadonaga, 2017). Also, like many early developmental genes, its precise pattern of expression in the blastoderm is coordinated by multiple enhancers (Figure 1A). These characteristics make the *kni* locus a good system in which to examine the roles of multiple promoters in a single gene locus.

**Figure 1.**
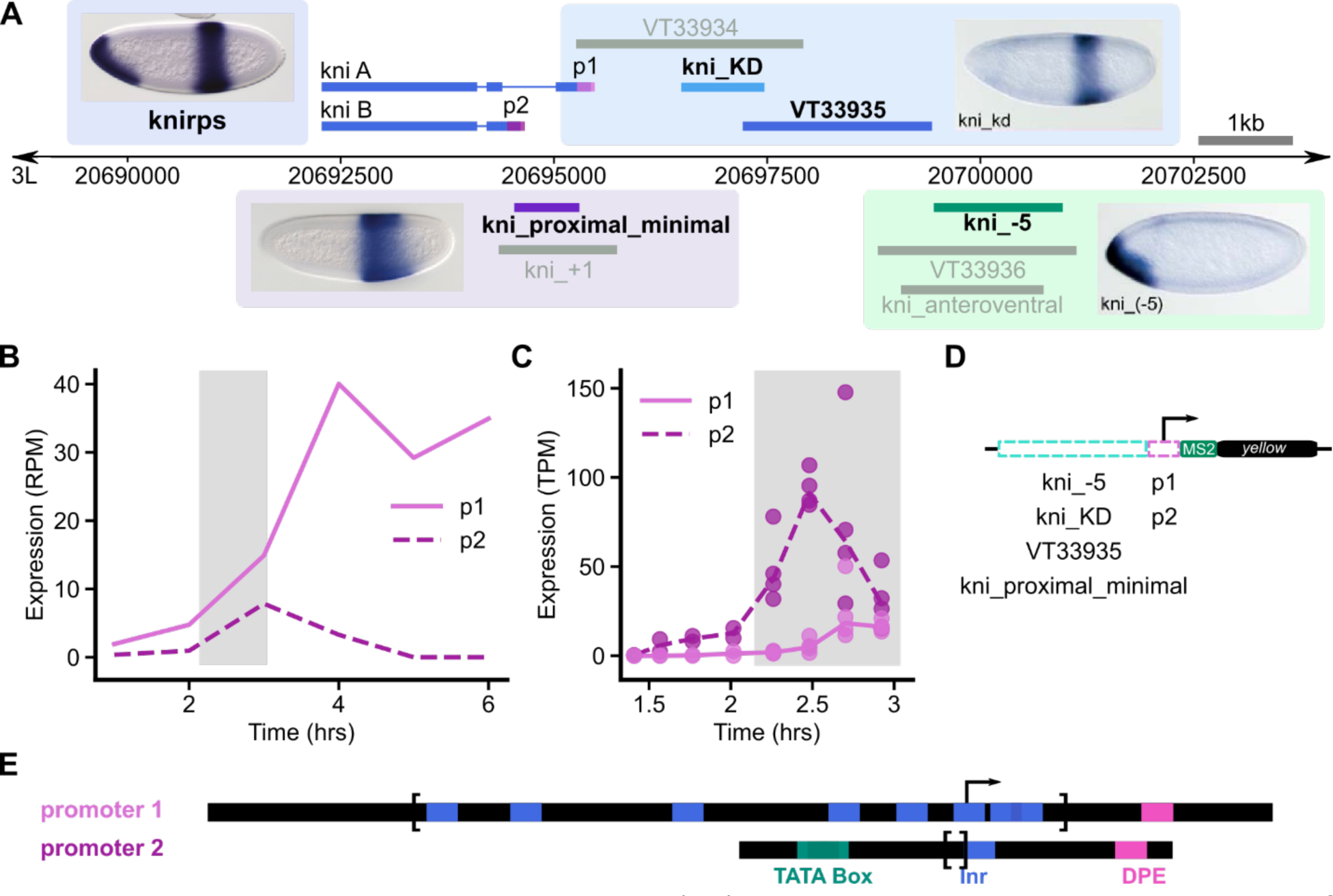
*knirps* as a case study. The *knirps* (*kni*) locus was chosen to study how the motif content of endogenous enhancers and promoters affects transcription dynamics. This locus was selected because it comprises multiple enhancers that bind different TFs and multiple core promoters that contain different promoter motifs. These enhancers and promoters are all active during the blastoderm stage. **(A)** The *kni* locus comprises multiple enhancers that together drive expression of a ventral, anterior band and a posterior stripe, as shown in the in situ at the top left. Enhancers that drive similar expression patterns have been displayed together in boxes with a representative *in situ* hybridization (Perry et al., 2011; Schroeder et al., 2004). The four enhancers selected for study are in color and labeled in bold text; the others are in gray. *kni* also has two promoters represented in two shades of purple, which drive slightly different transcripts (differing by only five amino acids). Expression data for the two *kni* promoters is shown, with RAMPAGE data (P. J. Batut & Gingeras, 2017) in **(B)** and RNA-seq data (Lott et al., 2011) in **(C)**; the time period corresponding to the blastoderm stage is highlighted in gray. Based on these two sets of data, the two *kni* promoters are both used during nuclear cycle 14 though which one is more active is less clear. Note that for the rest of development, promoter 1 is the more active one. **(D)** A total of eight MS2 reporter constructs containing pairs of each of the four enhancers matched with each of the two *kni* promoters were made. **(E)** The two *kni* promoters are shown here in black, consisting of the RAMPAGE-defined transcription start clusters (TSCs) between the brackets and an additional ± 40bp from the TSCs. The two *kni* promoters can be distinguished by their motif content (with promoter 1 consisting of a series of Inr motifs and a DPE motif and promoter 2 consisting of an Inr, two overlapping TATA Boxes and a DPE motif). They also differ in the “sharpness” of their region of transcription initiation (shown between the brackets), with promoter 1 (124bp) being significantly broader than promoter 2 (3bp) based on RAMPAGE tag data (P. J. Batut & Gingeras, 2017).

We used several approaches to delineate the roles of these two promoters. To examine the molecular compatibility of different *kni* enhancer-promoter pairs in a controlled setting, we created reporter constructs of eight *kni* enhancer-promoter pairs driving expression of an MS2 reporter. We found that some *kni* enhancers are able to interact with multiple promoters similarly, while others have a strong preference for one. By using the MS2 system to measure the transcription dynamics, we also determined the molecular events that lead to these preferences. Next, analysis of a *kni* locus reporter demonstrated that locus context can affect promoter-enhancer preferences and indicates that promoters both have different jobs and provide some amount of redundancy. Finally, we explored the role of different promoter motifs in specifying expression dynamics by using constructs with promoter mutations. Examining the *kni* locus has allowed us to (1) determine how transcription dynamics are impacted by molecular compatibility, (2) determine the roles of multiple promoters in a locus, and (3) probe how the motif content of promoters produces a particular expression output.

## Results

### Selection of enhancers and promoter pairs tested

*knirps* has two conserved promoters that drive very similar transcripts (Figure 1A; Figure S1A and B). Most previous studies discuss the role of a single *kni* promoter (promoter 1), though in practice, many of the constructs used in these studies actually contained both promoters, since promoter 2 is located in a *kni* intron (Bothma et al., 2015; El-Sherif & Levine, 2016; Pankratz et al., 1992; Pelegri & Lehmann, 1994). While more transcripts initiate from promoter 1 throughout most of development (Figure 1B), based on two different measures of transcript abundance, both promoters appear to be active during nuclear cycle 14, 2-3 hours after fertilization (Figure 1B and 1C) (P. J. Batut & Gingeras, 2017; Lott et al., 2011). These two promoters are distinguished by their motif content and by their “shape” (Figure 1E). Promoter 1 is composed of multiple Initiator (Inr) motifs, each of which can specify a transcription start site. These Inr motifs enable promoter 1 to drive transcription initiation in a 124 bp window, characteristic of a “broad” or “dispersed” promoter typically associated with housekeeping genes (Juven-Gershon, Hsu, Theisen, & Kadonaga, 2008; Sloutskin et al., 2015). There is a single DPE element in promoter 1; however, its significance is somewhat unclear, as it is only the canonical distance from a single, somewhat weak, Inr motif within the initiation window. Promoter 2 is composed of Inr, TATA Box and DPE motifs. This motif structure leads promoter 2 to initiate transcription in a 3 bp region, which is characteristic of the “sharp” or “focused” promoter shape typically associated with developmental genes (Figure S1C).

To select key early embryonic *kni* enhancers, we took into account the expression patterns driven by the enhancers and their overlap in the locus. We split the enhancers into three groups based on their expression patterns and selected one representative enhancer per group— enhancers driving a diffuse posterior stripe (kni_proximal_minimal), enhancers driving a sharp posterior stripe (kni_KD, the “classic” *kni* posterior stripe enhancer), and enhancers driving the anterior band (kni_-5) (Figure 1A). Among the enhancers driving a sharp posterior stripe, we decided to examine another enhancer, VT33935, in addition to kni_KD (Pankratz et al., 1992). VT33935 was identified in a high-throughput screen for enhancer activity (Kvon et al., 2014) and has only minimal overlap with the kni_KD enhancer but drives the same posterior stripe of expression. This suggests it may be an important contributor to *kni* regulation.

To determine the TF inputs to these enhancers, we scanned each enhancer using the motifs of TFs regulating early axis specification and calculated an overall binding capacity for each enhancer-TF pair (Figure 2A and S2). We found that kni_KD and VT33935 seem to be regulated by similar TFs, which suggests that together they comprise one larger enhancer. Here, we studied them separately, as historically kni_KD has been considered the canonical enhancer driving posterior stripe expression (Pankratz et al., 1992). Since kni_KD, VT33935, and kni_proximal_minimal drive overlapping expression patterns, they can be considered a set of shadow enhancers. Despite their similar expression output, kni_proximal_minimal has different TF inputs than the other two, including different repressors and autoregulation by Kni itself (Perry, Boettiger, & Levine, 2011). kni_-5 is the only enhancer that controls expression of a ventral, anterior band. Accordingly, this is the only enhancer of the four that has dorsal-ventral TF inputs (Dorsal and Twist) (Figure 2A) (Schroeder et al., 2004). In sum, analyses of the total binding capacity of these enhancers demonstrate that they are bound by different TFs (Figure 2A).

**Figure 2.**
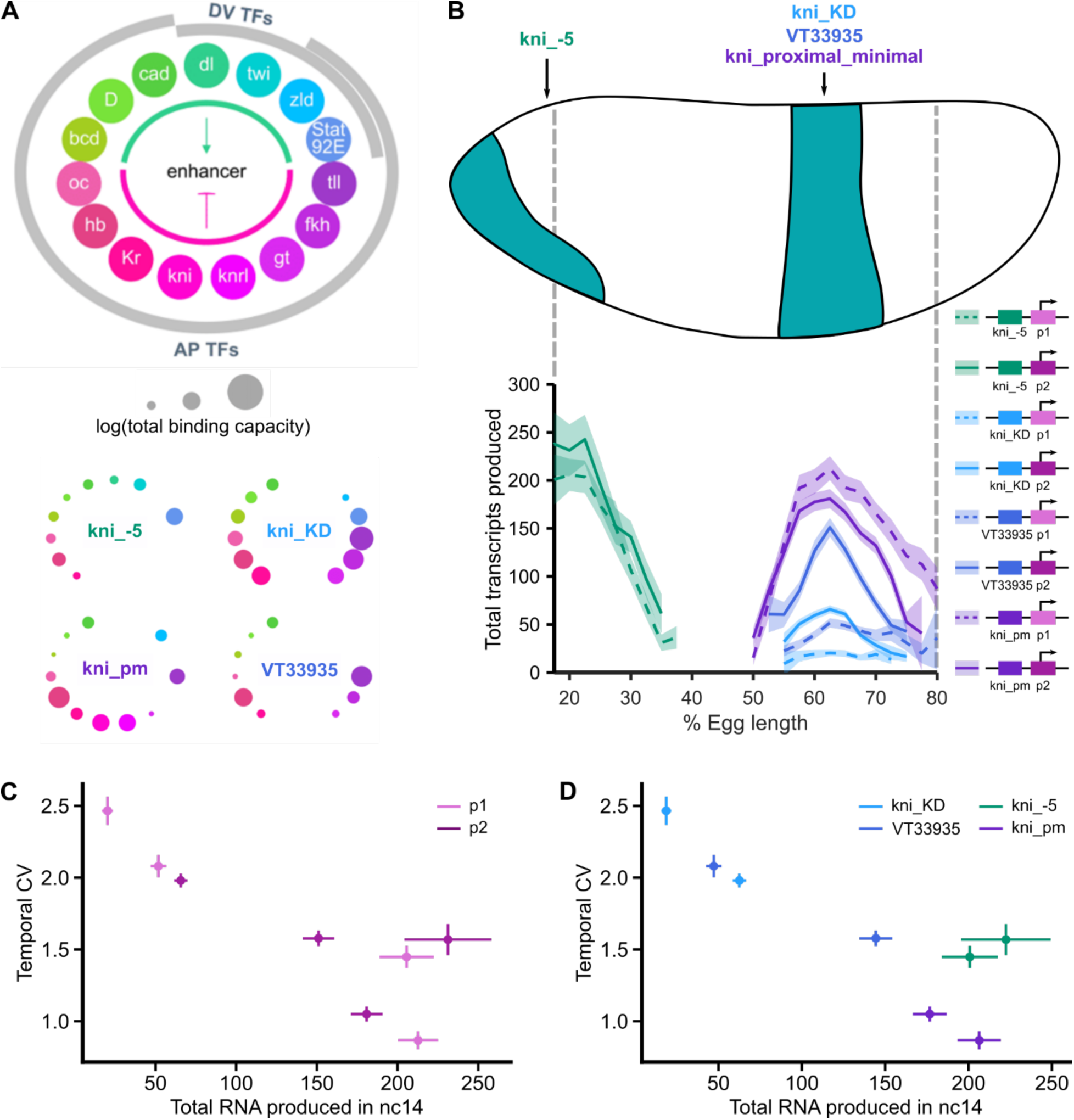
The *kni* enhancers differ in their capacity to bind different transcription factors and drive transcription with different promoters. The enhancers can be separated into two classes—those that produce high expression with either promoter (kni_-5 and kni_proximal_minimal) and those that produce much higher expression with promoter 2 (kni_KD and VT33935). Note that for simplicity, kni_proximal_minimal has been shortened to kni_pm in the figures. **(A)** Here ability of the *kni* enhancers to bind early axis-patterning TFs is quantified and represented visually. The logarithm of the predicted TF binding capacity of each of the *kni* enhancers is plotted as circles around the enhancer, with the color indicating the TF and the circle size increasing with higher binding capacity. The TFs are categorized by their role in regulating anterior-posterior (AP) or dorsal-ventral (DV) patterning and broadly by their roles as activators (indicated by the green arc) and repressors (indicated by the pink arc). Note that kni_KD and VT33935, which drive the same posterior stripe of expression, share very similar TFs and that kni_-5, the only enhancer with a DV component, is the only one bound by DV TFs. Kni_proximal_minimal drives a similar expression pattern to kni_KD and VT33935, but notably has different predicted TF binding capacities. **(B)** The *Drosophila* embryo with the *kni* expression pattern at nuclear cycle 14 is shown; kni_-5 drives the expression of the anterior, ventral band, while the other three enhancers drive the expression of the posterior stripe. We made enhancer-promoter reporters containing each of the four enhancers matched with either promoter 1 or 2. Using measurements from these enhancer-promoter reporters (shown at the right), the total RNA produced by each construct during nuclear cycle 14 is plotted against position along the embryo length (AP axis). The error bands around the lines are 95% confidence intervals. The constructs containing promoter 1 are denoted with a dashed line and those containing promoter 2 with a solid line. Some, but not all, enhancers show a strong promoter preference. kni_KD and VT33935, which are bound by similar TFs, drive 2.9-fold and 3.4-fold higher expression with promoter 2 at 62.5% embryo length, respectively (one-sided *t*-test *p* < 2.2 x 10^-16^ for both), whereas, kni_-5 and kni_proximal_minimal show similar expression regardless of promoter with the largest difference only 1.2-fold at the anterior-posterior bin of maximum expression (22% and 63%, respectively) (two-sided *t*-test comparing kni_-5-promoter1 vs. kni_-5-promoter2, *p* = 0.12 and kni_proximal_minimal-promoter1 vs. kni_proximal_minimal-promoter2 *p* = 9.8 × 10^-5^). In panels **(C – D)**, the temporal coefficient of variation (CV) is plotted against the total RNA produced in nc14 at the anterior-posterior bin of maximum expression (22% and 63%) for the anterior band and the posterior stripe, respectively, with the error bars representing 95% confidence intervals. There is a general trend of mean expression levels being anti-correlated with CV, or noise. **(C)** Here, the data points are colored by the construct’s promoter, with promoter 1 in light purple and promoter 2 in purple. Despite the general trend, there are cases when the same promoter (promoter 2) shows higher CV and total expression when paired with different enhancers (kni_proximal_minimal vs kni_-5). **(D)** Here, the data points are colored by the construct’s enhancer. Again, despite the general trend, there are cases when the same enhancer (kni_-5) shows higher CV and total expression when paired with different promoters (promoter 1 vs 2).

By using this set of endogenously interacting enhancers and promoters with varied motif content, we can elucidate the functional value of having multiple promoters. In particular, we can determine whether multiple promoters exist because different enhancers work with different promoters, or whether having multiple promoters provides necessary redundancy in the system, or some combination of the two.

### Some enhancers tolerate promoters of different shapes and composition

To characterize the inherent ability of promoters and enhancers to drive expression, without complicating factors like enhancer competition, promoter competition, or variable enhancer-promoter distances, we created a series of eight transgenic enhancer-promoter reporter lines. Each reporter contains one enhancer and one promoter directly adjacent to each other, followed by MS2 stem loops inserted in the 5’ UTR of the *yellow* gene (Figure 1D, see Methods for details). These tagged transcripts are bound by MCP-GFP fusion proteins, yielding fluorescent puncta at the site of nascent transcription. The fluorescence intensity of each spot is proportional to the number of transcripts in production at a given moment (Garcia, Tikhonov, Lin, & Gregor, 2013).

When considering the expression output driven by these enhancer-promoter combinations, several outcomes are possible. One possible outcome is that one promoter is simply stronger than the other – consistently driving higher expression, regardless of which enhancer it is paired with. Another possibility is that each enhancer drives higher expression with one promoter than with the other, but this preferred promoter differs between enhancers. This would suggest that promoter motifs and shape affect their ability to successfully interact with enhancers with different bound TFs to drive expression. Lastly, it is possible that some enhancers drive similar expression with either promoter, this suggests that the particular set (and orientation) of the TFs recruited to those enhancers allow them to transcend the differences in promoter architecture.

When comparing the mean expression levels, we found that some enhancers (kni_-5 and kni_proximal_minimal) have relatively mild preferences for one promoter over the other (Figure 2B; two-sided *t*-test comparing kni_-5-promoter1 vs. kni_-5-promoter2, *p* = 0.12 and kni_proximal_minimal-promoter1 vs. kni_proximal_minimal-promoter2 *p* = 9.8 × 10^-5^). Despite the significant differences between these enhancer-promoter constructs, the effect size is relatively small, with the largest difference in mean expression being 1.2-fold. This suggests that the TFs recruited to these enhancers can interact with very different promoters more or less equally well. On the other hand, kni_KD and VT33935 respectively drive 2.9-fold and 3.2-fold higher expression with promoter 2 than promoter 1 at 62.5% embryo length (Figure 2C; one-sided *t*-test *p* < 2.2 x 10^-16^ for both). This suggests that the TFs recruited to kni_KD and VT33935, which are similar, (Figure 2A) limit their ability to successfully drive expression with promoter 1, which is a dispersed promoter. Taken together, this implies a simple model of promoter strength is not sufficient to account for these results. Instead, it is the combination of the proteins recruited to both enhancers and promoters that set expression levels, with some enhancers interacting equally well with both promoters and others having a preference.

These differences in enhancer preference or lack thereof may be mediated by the particular TFs recruited to them and the motifs present in the promoters. Previous researchers have found that the developmental TFs, Caudal (Cad) and Dorsal (Dl), tend to regulate genes with DPE motifs and drive lower expression when DPE has been eliminated (Juven-Gershon, Hsu, & Kadonaga, 2008; Zehavi et al., 2014). In addition, computational analysis of TF-promoter motif co-occurrence patterns indicates that Bcd shows a similar enrichment for DPE-containing promoters and a depletion for Inr- and TATA box-containing promoters when DPE is absent (Figure S2). A study also indicated that Bcd can work in conjunction with Zelda to activate a TATA Box-containing promoter, but this combination does not appear to be widely generalizable (Ling et al., 2019). In accordance with that, we find that all four *kni* enhancers, which bind Cad and Bcd, drive relatively high expression with the DPE-containing promoter 2. Interestingly, in the case of kni_-5 and kni_pm, we find that they can also drive similarly high expression with the series of weak Inr sites that composes promoter 1. This indicates that while some factors mediating enhancer-promoter preference have been identified, there are additional factors we have yet to discover that are playing a role.

We also calculated the expression noise associated with each construct and plotted it against the expression output of each. Previous studies have suggested that TATA-containing promoters generally drive more noisy expression (Ramalingam, Natarajan, Johnston, & Zeitlinger, 2021; Ravarani, Chalancon, Breker, de Groot, & Babu, 2016). Among our constructs, expression noise is generally inversely correlated with mean expression (Figure 2C and 2D), and the TATA-containing promoter 2 does not have uniformly higher noise than the TATA-less promoter 1. However, some constructs, notably those containing kni_-5, have higher noise than others with similar output levels, suggesting that, in this case, promoters alone do not determine expression noise.

### Simple model of transcription and molecular basis of burst properties

To unravel the molecular events that result in these expression differences, we consider our results in the context of the two-state model of transcription (Peccoud & Ycart, 1995; Tunnacliffe & Chubb, 2020). Here, the promoter is either (1) in the inactive state (“OFF”), in which RNA polymerase cannot initiate transcription or (2) in the active state (“ON), in which it can (Figure 3A). The promoter transitions between these two states with rates *k_on_* and *k_off_*, with the transitions involving both the interaction of the enhancer and promoter and the assembly of the necessary transcriptional machinery. This interaction may be through direct enhancer-promoter looping or through the formation of a transcriptional hub, a nuclear region with a high concentration of TFs, co-factors, and RNA polymerase (Lim & Levine, 2021). For simplicity, we will use looping as a shorthand to include both scenarios. In its active state, the promoter produces mRNA at rate *r*, and given our ability to observe only nascent transcripts, the mRNA decay rate *μ* denotes the diffusion of mRNA away from the gene locus.

**Figure 3.**
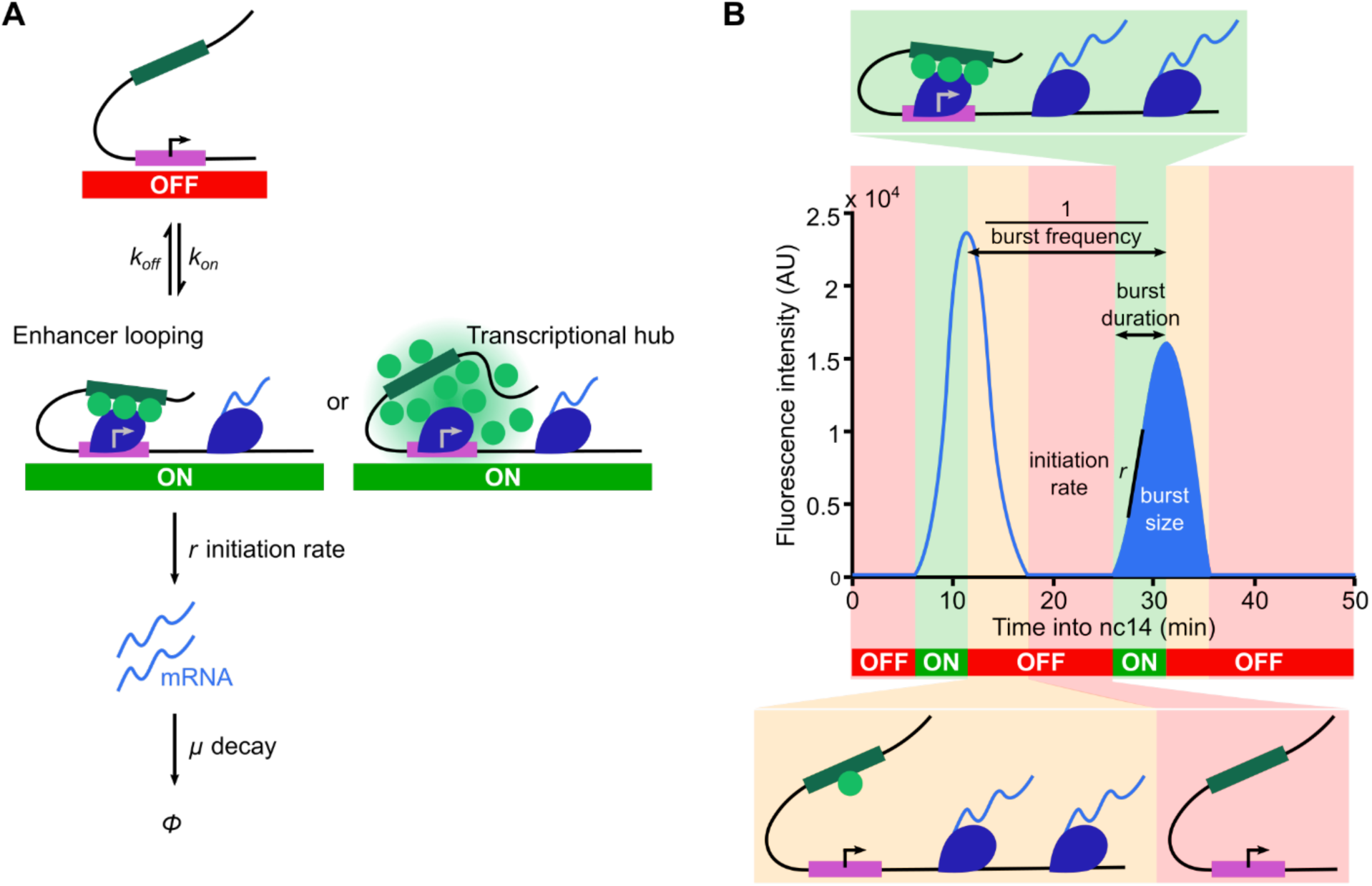
Two-state model of transcription in the context of tracking transcription dynamics. **(A)** Here, we represent the two-state model of transcription, in which the promoter is either (1) in the inactive state (OFF), in which RNA polymerase cannot bind and initiate transcription or (2) in the active state (ON), during which it can. The promoter transitions between these two states with rates *k_on_* and *k_off_*, with promoter activation involving both the interaction of the enhancer and promoter and the assembly of all the necessary transcription machinery for transcription initiation to occur. This may occur through enhancer looping or through the formation of a transcriptional hub. In its active state, the promoter produces mRNA at rate *r*, and the mRNA decays by diffusing away from the gene locus at rate *μ*. **(B)** MS2-tagging RNA allows us to track nascent transcription, and the resulting fluorescence trace (in light blue) is proportional to the number of nascent RNA produced over time. The graph is split into sections, representing different molecular states and how they correspond to fluctuating transcription over time. These states are represented by different colors—red when the promoter is OFF, green when it is ON, and yellow when transcription continues but the promoter is no longer ON, as no new polymerases are being loaded. The dynamics of these fluctuations or bursts can be characterized by quantifying various properties, including burst frequency (how often a burst a occurs), burst size (number of RNA produced per burst), and burst duration (the period of active transcription during which mRNA is produced at rate *r*).

We track these molecular events by analyzing the transcription dynamics driven by each reporter and quantifying several properties. Total expression is simply the integrated signal driven by each reporter. The burst duration is the period of active transcription, and is dependent on *k_off_*, the rate of dissociation of enhancer and promoter looping (Figure 3B). The burst size, or number of transcripts produced per burst, depends on the burst duration and the RNA Pol II initiation rate. (Short, aborted transcripts and paused PolII are not visible in MS2 measurements). The burst frequency, or the inverse of the time between two bursts, depends on both *k_on_* and *k_off_*. Previous work in the early embryo has shown burst duration (and thus *k_off_*) to be reasonably consistent regardless of enhancer and promoter (Waymack, Fletcher, Enciso, & Wunderlich, 2020; Yokoshi, Cambón, & Fukaya, 2021). Within this regime, burst frequency is mainly dependent on *k_on_*. We used this model to characterize how the transcription output produced is affected by different combinations of the *kni* enhancers and promoters.

### Using GLMs to parse the role of enhancers, promoters, and their interactions

To parse the role of enhancers, promoters, and their interactions more clearly in determining expression levels in these reporters, we built separate generalized linear models (GLMs) to describe each transcriptional property. We visually represented the model using a bar graph (Figure 4A) in which the contributions of enhancer, promoter, and their interactions are represented in bars of green, purple, and brown, respectively (Figure 4B). Since the relative differences in expression driven by different enhancer-promoter pairs are generally consistent across the AP axis, we used the expression levels at the location of maximum expression along the AP axis (22% and 63% for the anterior band and posterior stripe, respectively, Figure 2C).

**Figure 4.**
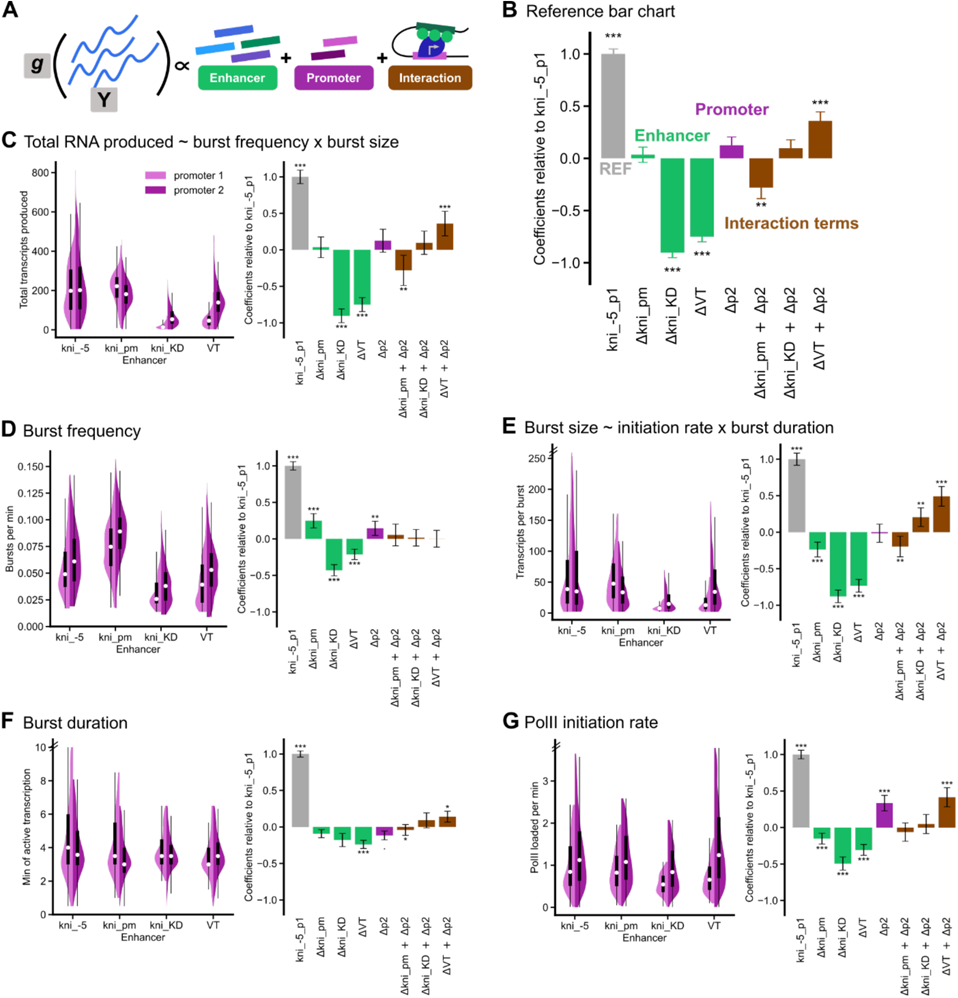
Expression levels are mainly determined by burst frequency and initiation rate. **(A)** To parse the effects of the enhancer, the promoter, and their interactions on all burst properties, we built generalized linear models (GLMs). Y represents the burst property under study, g is the identity link function, and the enhancers, promoters, and their interaction terms are the explanatory variables. The coefficients of each of these explanatory variables is representative of that variable’s contribution to the total value of the burst property. **(B)** All burst property data was taken from the anterior-posterior bin of maximum expression (22% and 63%) for the anterior band and the posterior stripe, respectively. The coefficients and the 95% confidence intervals for each independent variable relative to that of a reference construct (kni_-5-p1) are plotted as a bar graph; * *p* < 0.05, ** *p* < 0.01, *** *p* < 0.001. The reference construct is represented in gray, and the effects of enhancer, promoter, and their interactions are represented in green, purple, and brown, respectively. Summing the relevant coefficients gives you the average value of the burst property for a particular construct relative to the reference construct. Thus, as the reference construct, kni_-5-p1 coefficient will always be 1. The average value of the burst property for a particular construct, e.g. VT-p2, relative to the reference construct, would be 0.75, which is the sum of the reference bar = 1, ΔVT = −0.78, Δp2 = 0.17, and ΔVT + Δp2 = 0.36. Note that for simplicity, kni_proximal_minimal and VT33935 has been shortened to kni_pm and VT, respectively, in the following graphs. In panels **(C – G)**, (left) split violin plots (and their associated box plots) of burst properties for all eight constructs will be plotted with promoter 1 in light purple and promoter 2 in purple. The black boxes span the lower to upper quartiles, with the white dot within the box indicating the median. Whiskers extend to 1.5*IQR (interquartile range) ± the upper and lower quartile, respectively. (right) Bar graphs representing the relative contributions of enhancer, promoter, and their interactions to each burst property are plotted as described in **(B)**. The double hash marks on the axes indicate that 90% of the data is being shown. **(C)** Expression levels are mainly determined by the enhancer and the interaction terms. Some enhancers (kni_-5 and kni_proximal_minimal) appear to work well with both promoters; whereas, kni_KD and VT, which are bound by similar TFs, show much higher expression with promoter 2. **(D)** Burst frequency is dominated by the enhancer and promoter terms, with promoter 2 consistently producing higher burst frequencies regardless of enhancer. **(E)** Burst size, which is determined by both initiation rate and burst duration, is dominated by the enhancer and interaction terms, with interaction terms representing the role of molecular compatibility. As **(F)** burst duration is reasonably consistent regardless of enhancer or promoter, differences in burst size are mainly dependent on differences in **(G)** PolII initiation rate.

If the molecular compatibility of the proteins recruited to the enhancer and promoter are important in determining a particular property, then we should find the interaction terms (in brown) to be sizeable in comparison with those of the enhancers (in green) and promoter (in purple). If not, the interaction terms will be relatively small. To develop an intuition for this formalism, we first built a GLM to describe total expression output. Using the GLM, we can see that enhancer, promoter, and interactions terms each play an important role in determining the expression output (Figure 4C), consistent with our qualitative interpretation above.

To determine which molecular events are modulated by molecular compatibility, we then applied this same GLM structure to each burst property. For example, molecular compatibility could increase the probability of enhancer-promoter loop formation, hence increasing the burst frequency. Alternatively, molecular compatibility could increase the rate at which RNA PolII initiates transcription, increasing burst size.

### Burst frequency and initiation rate are the primary determinants of expression levels

We found that the differences in total expression output are primarily mediated through differences in burst size (Figure 4E) and burst frequency (Figure 4D). Burst duration is very consistent across all constructs (Figure 4F). While the enhancer, promoter, and interaction terms all have a significant impact on duration (multivariate ANOVA; enhancer: *p* = 4.4 × 10^-10^; promoter: *p* = 4.1 × 10^-5^; interaction: *p* = 4.6 × 10^-5^), the effect size is small, with the largest difference being only 1.3-fold. Since burst size can be modulated by initiation rate and burst duration, and burst duration is relatively constant, this suggests that initiation rate and burst frequency are the primary dials used to tune transcription in these synthetic constructs.

Burst size is strongly dependent on both the enhancer and interaction terms; the interaction terms are a proxy for molecular compatibility. Of the variability in burst size explained by this model, enhancers and interaction terms account for 67.6% and 23.7% of the variance, respectively (Figure 4E). The differences in burst size were mainly achieved by tuning PolII initiation rate (Figure 4G). Conversely, burst frequency is dependent on promoter and enhancer identity, with negligible interaction terms (Figure 4D). Since burst frequency mainly depends on association rate (*k_on_*), this suggests that both enhancers and promoters play a large role in determining the likelihood of promoter activation, with molecular compatibility only minimally affecting this likelihood.

It is somewhat surprising that molecular compatibility plays only a small role in determining *k_on_*, since one might expect the interactions between the proteins recruited to promoters and enhancers would determine the likelihood of promoter-enhancer looping. This may be the result of the design of these constructs, with promoters and enhancer immediately adjacent to each other, and this may differ in a more natural context (see below). However, we do observe that molecular compatibility is important in determining the PolII initiation rate. This suggests that the TFs and cofactors recruited to each reporter may act synergistically to both recruit RNA PolII to the promoter and promote its successful initiation. In sum, these results indicate that not only do enhancer, promoter, and their molecular compatibility affect expression output, but they do so by tuning different burst properties in this synthetic setting.

### Despite promoter 2’s compatibility with kni_-5, promoter 1 primarily drives anterior expression in the locus context

The constructs measured thus far only contain a single enhancer and promoter, and therefore measure the inherent ability of a promoter and enhancer to drive expression. However, in the native locus, other complications like differing enhancer-promoter distances, enhancer competition, or promoter competition may impact expression output. To measure the effect of these complicating factors, we cloned the entire *kni* locus into a reporter construct and measured the expression patterns and dynamics of the wildtype locus reporter (wt) and reporters with either promoter 1 or 2 knocked out (Δp1 and Δp2) (Figure 5A). Due to the large number of Inr motifs, we made the Δp1 construct by replacing promoter 1 with a piece of lambda phage DNA. To make the Δp2 construct, we inactivated the TATA, Inr, and DPE motifs by making several mutations (see Methods for additional details).

**Figure 5.**
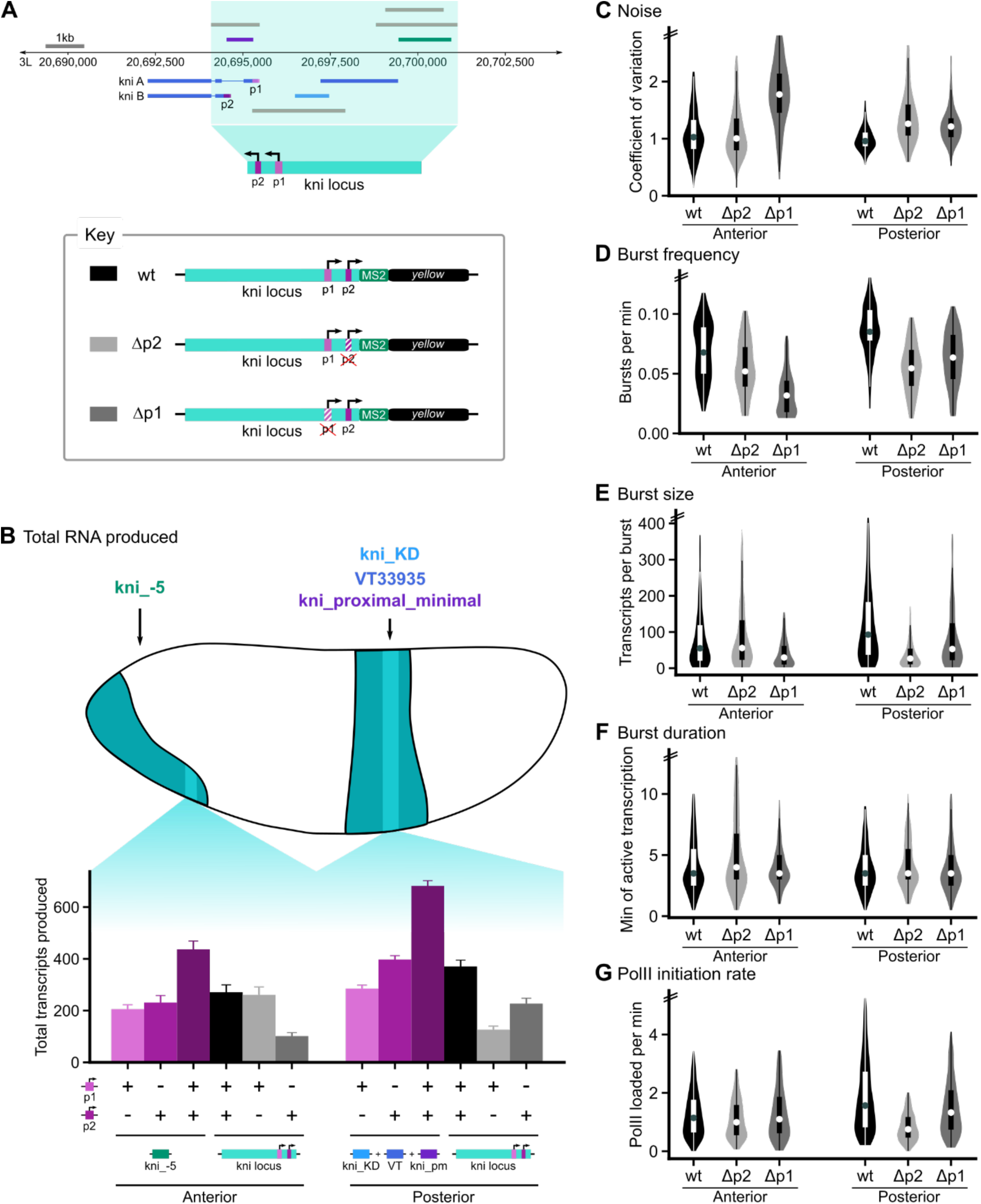
The synthetic enhancer-promoter constructs are insufficient to capture the behavior of the *knirps* promoters within the endogenous locus. **(A)** We cloned the entire *kni* locus into an MS2 reporter construct and measured the expression levels and dynamics of the wildtype (wt) locus reporter, and reporters with either promoter 1 or 2 knocked out (Δp1 and Δp2). To make the Δp1 reporter, we replaced promoter 1 with a piece of lambda phage DNA, due to the large number of Inr motifs. To make the Δp2 construct, we removed the TATA, Inr, and DPE motifs by making several mutations (see Methods for additional details). In panels **(B – G)**, all burst property data was taken from the anterior-posterior bin of maximum expression (22% and 63%) for the anterior band and the posterior stripe, respectively. **(B)** The *Drosophila* embryo with the *kni* expression pattern at nuclear cycle 14 is shown; kni_-5 drives the expression of the anterior, ventral band, while the other three enhancers drive the expression of the posterior stripe. The bin of maximum expression is highlighted in light teal. To compare the expression produced by the synthetic enhancer-promoter reporters with the locus reporters, we plotted bar graphs of the summed total RNA produced at the location of maximum expression in the anterior (left) and posterior (right) for six cases—just enhancer-promoter1 reporters (light purple), just enhancer-promoter2 reporters (purple), both enhancer-promoter1 and -promoter2 reporters (dark purple), the wt locus reporter (black), the locus Δp2 reporter (light gray), and the locus Δp1 reporter (dark gray). In panels **(C - F)** violin plots (and their associated box plots) of burst properties for all three reporters are plotted with the wt, Δp1, and Δp2 reporters in black, light gray, and dark gray, respectively. The internal boxes span the lower to upper quartiles, with the dot within the box indicating the median. Whiskers extend to 1.5*IQR (interquartile range) ± the upper and lower quartile, respectively. The double hash marks on the axes indicate that 95% of the data is being shown. **(C)** The coefficient of variation is inversely correlated with total RNA produced shown in **(B)**. In the anterior, the Δp2 reporter, which produces the same total RNA as the wt reporter, also produces the same amount of noise. **(D)** In the anterior of the embryo, burst frequency of the Δp2 reporter is less than the wt reporter even though they produce the same expression levels and noise. In the posterior, knocking out promoter 2 has a larger impact on burst frequency than knocking out promoter 1. **(E)** In both the anterior and posterior, burst size is directly correlated with total RNA produced. Note that in the posterior of the embryo, knocking out promoter 2 has a much larger impact on burst size than knocking out promoter 1. Burst size is dependent on PolII initiation rate and burst duration. While **(F)** burst duration is reasonably consistent regardless of promoter knockout, **(G)** PolII initiation rate is directly correlated with burst size. This suggests that differences in burst size are mainly mediated by differences in PolII initiation rate.

In the anterior, the kni_-5 enhancer is solely responsible for driving expression. Therefore, by comparing the expression output from the wildtype locus reporter and the kni_-5-promoter reporters in the anterior, we can measure the effect of the locus context, i.e. multiple promoters, differing promoter-enhancer distance, or other DNA sequence features. If the kni_-5-promoter reporters capture their ability to drive expression in the locus context, we would expect the locus reporter to drive expression equal to the sum of the kni_-5-p1 and kni_-5-p2 reporters. In contrast to this expectation, in the anterior band, the locus reporter drives a much lower level of expression than the sum of the two kni_-5 reporters (Figure 5B, dark purple vs black bar). In fact, the level is similar to the expression output of kni_-5 paired with either individual promoter, suggesting that kni_-5’s expression output is altered by the locus context.

The observed sub-additive behavior may arise in several ways. It may be that promoter competition similarly reduces the expression output of both p1 and p2 in the anterior. In this case, knocking out either promoter would produce wildtype levels of expression, as competition would be eliminated. Alternatively, the ability to drive expression in the locus context could be uneven between the promoters. If this is the case, we would expect the promoter knockouts to have different effects on expression.

Consistent with the second scenario, we find that when promoter 2 is eliminated in the *kni* locus construct, the expression in the anterior remains essentially the same (two-sided *t*-test comparing mean expression levels of wt vs. Δp2, *p* = 0.62), while a promoter 1 knockout has a significant impact on expression levels (one-sided *t*-test comparing mean expression levels of wt vs. Δp1, *p* < 2.2 × 10^-16^; Figure 5B). Thus, promoter 1 is sufficient to produce wildtype expression levels and patterns in the locus. The noise and the burst properties of the WT *kni* locus construct and the promoter 2 knockout are also nearly identical to the wildtype locus, further supporting the claim of promoter 1 sufficiency in the anterior (Figure 5C – G). Notably, even in a locus that contains promoters with and without a canonically placed DPE element (promoter 2 vs promoter 1), a Cad- and Dl-binding enhancer like kni_-5, can still primarily rely on the DPE-less promoter 1 to drive transcription.

When promoter 1 is eliminated from the locus, expression is cut to about one third of that of the wildtype locus construct, which is also lower than the expression output of the kni_-5-p2 construct. Thus, unlike promoter 1, promoter 2 loses its ability to drive wildtype levels of expression in the context of the locus. As promoter 2 is ∼650bp upstream of promoter 1, this extra distance between kni_-5 and promoter 2 may be sufficient to reduce promoter 2’s ability to drive expression. Alternatively, other features of the *kni* locus, such as the binding of other proteins or topological constraints, may interfere with the ability of the kni_-5 enhancer to effectively interact with promoter 2. The drop in expression is mediated by a tuning down of all burst properties (Figure 5D – G). In sum, the kni_-5 enhancer preferentially drives expression via promoter 1 in the locus, even though enhancer-promoter constructs indicate that it is equally capable of driving expression with promoter 2. When promoter 1 is absent from the locus, promoter 2 is able to drive a smaller amount of expression, suggesting that it can serve as a backup, albeit an imperfect one.

### In the posterior, both promoters are required for wildtype expression levels

The posterior stripe is controlled by three enhancers, with kni_proximal_minimal producing similar levels of transcription with either promoter, and the other two enhancers strongly preferring promoter 2 and driving lower expression overall (Figure 2B). Therefore, when considering the posterior stripe, the expression output of the locus reporter may differ from the individual enhancer-reporter constructs due to promoter competition, enhancer competition, different promoter-enhancer distances, or other DNA features. By comparing the sum of the six relevant enhancer-promoter reporters to the output of the locus reporter, we can see that the locus construct drives considerably lower expression levels than the additive prediction (Figure 5B, dark purple vs black bar). In fact, the locus reporter output levels are similar to the sum of the enhancer-promoter 2 reporters, suggesting that promoter 2 could be solely responsible for expression in the posterior, despite kni_proximal_minimal’s ability to effectively drive expression with promoter 1. If promoter 2 is sufficient for posterior stripe expression, we would predict that the promoter 1 knockout would have a relatively small effect, while a promoter 2 knockout would greatly decrease expression in the posterior.

In contrast to this expectation, both promoter 1 and promoter 2 knockouts have a sizable effect on expression output, indicating that both are required for wildtype expression levels in the posterior (Figure 5B, light gray and gray bars). Specifically, knocking out promoter 2 severely reduces expression in the posterior stripe, producing about half the expression of the summed outputs of the enhancer-promoter1 constructs (Figure 5B, light gray vs light purple bars). Knocking out promoter 1 also reduces expression in the posterior stripe but not as severely as knocking out promoter 2 (Figure 5B, gray vs light gray bars). The promoter 1 knockout generates about half the expression of the summed expression output of the enhancer-promoter2 constructs (Figure 5B, gray vs purple bars). In both cases, the results indicate that the differences in locus context cause the enhancers to act sub-additively, even when only one promoter is present.

The promoter knockouts also allow us to examine how they tune expression output. Knocking out either promoter impacts burst size (and thus initiation rate) and burst frequency, though knocking out promoter 2 has a more severe impact (Figure 5D, 5E and 5G). These results show that, in the posterior, both promoters are required to produce WT expression levels when considered in the endogenous locus setting (Figure 5B, light and dark gray vs black bars). This is despite the fact that enhancer-promoter reporters indicate that, in the absence of competition, promoter 2 alone would suffice (Figure 5B, purple vs black bars).

### PolII initiation rate is a key burst property that is tuned by promoter motif

Studying these enhancers and promoters in the locus context demonstrated that distance and competition affect a promoter’s ability to drive expression, but now we narrow our focus to promoter 2’s remarkable compatibility with enhancers that bind very different sets of TFs. To dissect how its promoter motifs enable promoter 2 to be so broadly compatible, we again made enhancer-promoter reporter lines in which one enhancer and one promoter are directly adjacent to each other, but this time the promoter is a mutated promoter 2 in which the TATA Box and DPE motifs have been eliminated (Figure 6A, see Methods for details). This allows us to determine whether a single, strong Inr site (mutated promoter 2) can perform similarly to a series of weak Inr sites (promoter 1) and to clarify the role of TATA Box and DPE motifs in tuning burst properties.

**Figure 6.**
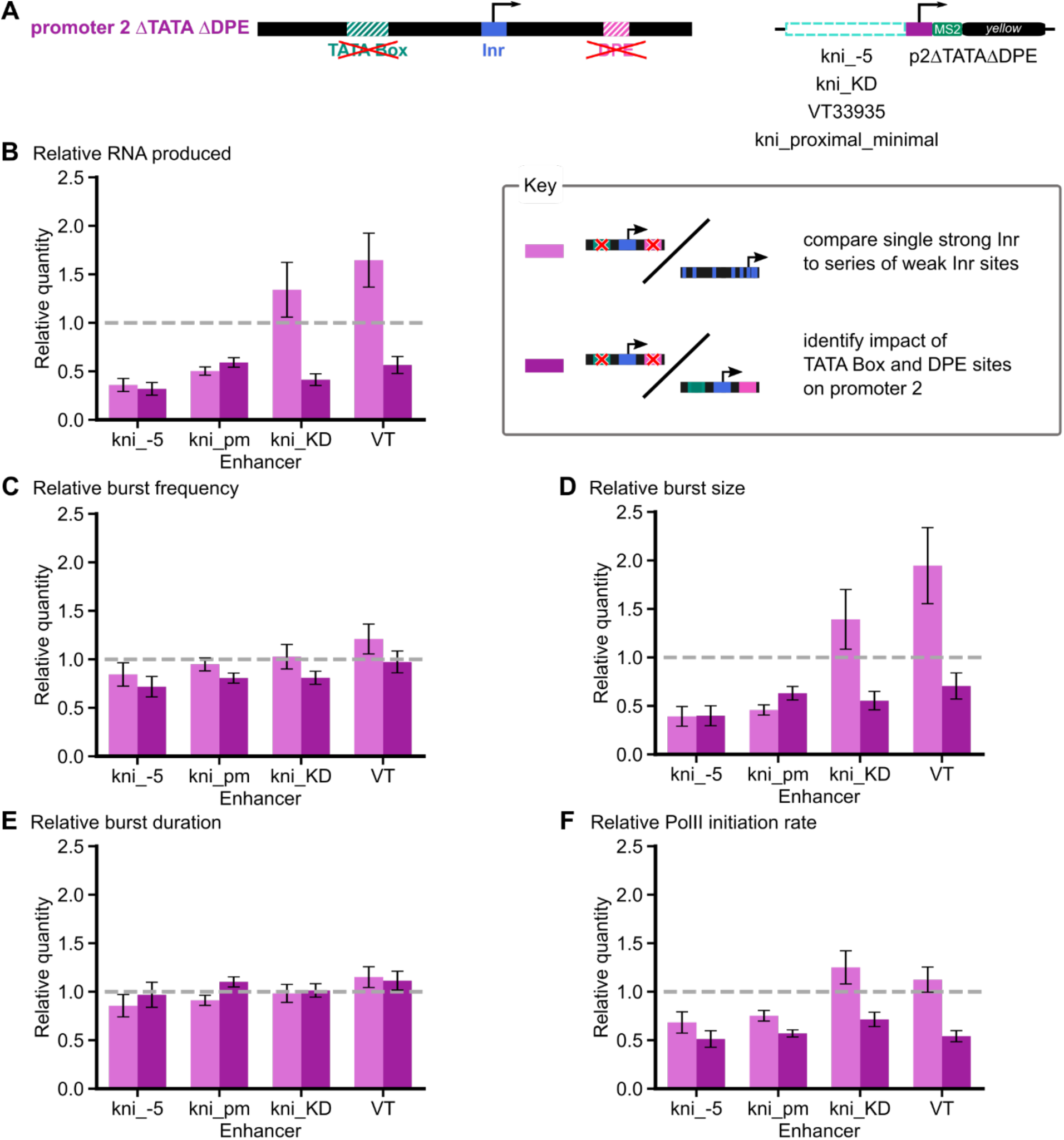
PolII initiation rate is a key burst property that is tuned by promoter motif. **(A)** We made enhancer-promoter reporters containing each of the four enhancers matched with a mutated promoter 2 (p2ΔTATAΔDPE) in which the TATA Box and DPE motifs have been eliminated by making several mutations (see Methods for details). In panels **(B – F)**, bar graphs of the burst properties produced by p2ΔTATAΔDPE relative to promoter 1 (in light purple) and to promoter 2 (in purple) are shown. By comparing p2ΔTATAΔDPE with promoter 1, we can determine whether a single, strong Inr site (mutated promoter 2) can perform similarly to a series of weak Inr sites (promoter 1), and by comparing p2ΔTATAΔDPE with promoter 2, we can clarify the role of TATA Box and DPE motifs in tuning burst properties. The error bars show the 95% confidence intervals. The gray dashed line at 1 acts as a reference—if there is no difference between the burst properties produced by p2ΔTATAΔDPE and either promoter 1 or 2, the bar should reach this line. All burst property data was taken from the anterior-posterior bin of maximum expression (22% and 63%) for the anterior band and the posterior stripe, respectively. Note that for simplicity, kni_proximal_minimal and VT33935 have been shortened to kni_pm and VT, respectively, in the following graphs. **(B)** When comparing p2ΔTATAΔDPE with promoter 1, we can see that the enhancers fall into two classes—those that drive less expression or more expression with a single strong Inr site than with a series of weak Inr sites. The enhancers (kni_-5 and kni_proximal_minimal) that drive less expression are the same ones that were similarly compatible with both promoters 1 and 2, whereas the enhancers that drive more expression (kni_KD and VT33935) are the ones that strongly preferred promoter 2. When comparing p2ΔTATAΔDPE with promoter 2, we see that eliminating TATA Box and DPE motifs reduces expression output for all enhancers. **(C)** When comparing p2ΔTATAΔDPE with either promoter 1 or promoter 2, we see that burst frequency is not substantially affected though, compared to promoter 2, there is a moderate decrease upon motif disruption. **(D)** When comparing the burst size of p2ΔTATAΔDPE reporters with either that of promoter 1 or promoter 2 reporters, we see the same behavior as with total RNA (shown in panel B). This suggests that burst size is the main mediator of the increase or decrease in total RNA produced. Burst size is dependent on PolII initiation rate and burst duration. As **(E)** burst duration is reasonably consistent regardless of promoter, it appears that **(F)** changes in burst size are mainly mediated by tuning PolII initiation rate. Together, this suggests that enhancers fall into two classes, based on their response to different promoters; however, regardless of class, PolII initiation rate is what underlies differences in expression output.

Promoter 2 is characterized by two TATA Boxes, an Inr motif, and a DPE motif. Previously, much research has focused on comparing TATA-dependent with DPE-dependent promoters; however, many promoters contain both. Here, we consider how the presence of both may impact transcription. We know that each of these motifs recruits subunits of TFIID, with TATA Box recruiting TBP or TRF1 (Hansen, Takada, Jacobson, Lis, & Tjian, 1997; Holmes & Tjian, 2000; Kim, Nikolov, & Burley, 1993), Inr recruiting TAF1 and 2 (Chalkley & Verrijzer, 1999; Wu et al., 2001), and DPE recruiting TAF6 and 9 (Shao et al., 2005), as well as other co-factors like CK2 and Mot1 (Hsu et al., 2008; Lewis, Sims, Lane, & Reinberg, 2005). Strict spacing between TATA-Inr and Inr-DPE both facilitate assembly of all these factors and others into a pre-initiation complex (Burke & Kadonaga, 1996; Emami, Jain, & Smale, 1997). It is likely that a promoter with all three motifs will behave similarly, with the addition of each motif further tuning the composition, configuration, or flexibility of the transcriptional complex. Given this, elimination of the TATA Box and DPE motifs may weaken the promoter severely through loss of cooperative interactions, especially for kni_KD and VT33935, which are significantly more compatible with promoter 2 than promoter 1. Alternatively, the single strong Inr site may be sufficient to recruit the necessary transcription machinery, especially in the case of kni_-5 and kni_proximal_minimal, which work well with the series of weak Inr sites that composes promoter 1.

When compared to promoter 1, we see that promoter 1-compatible enhancers (kni_-5 and kni_proximal_minimal) drive lower expression with a single Inr than with a series of weak Inr sites (Figure 6B, light purple bars). In contrast, enhancers less compatible with promoter 1 (kni_KD and VT33935) drive higher expression with a single Inr site than promoter 1 even without the TATA Box and DPE sites (Figure 6B, light purple bars), suggesting that the strong Inr is the key to better expression output with these enhancers. For all enhancers, the resulting expression change appears to be mediated mainly through a decrease in burst size due to a reduction in initiation rates (Figure 6D – F).

Given that all four enhancers are compatible with promoter 2, and promoter 2 appears to achieve higher expression by tuning PolII initiation rates, we posit that TATA Box and DPE are what help promoter 2 drive high initiation rates. When comparing p2ΔTATAΔDPE with promoter 2, we see that all enhancers produce lower expression (Figure 6B, dark purple bars), and this is mediated mainly through tuning burst size (Figure 6D) and, for some enhancers, also burst frequency (Figure 6C). Notably, burst size (and thus polymerase initiation rate), which were most dependent on molecular compatibility, are affected the most by the elimination of the TATA Box and DPE motifs (Figure 6D and 6E), indicating that molecular compatibility plays an important role mediating high expression output. Interestingly, even in the absence of the TATA Box and DPE motifs, the one strong Inr site is sufficient to produce higher expression with the enhancers less compatible with promoter 1 (kni_KD and VT33935), and this increased expression is also mediated by higher polymerase initiation rates (Figure 6B and 6F, light purple bars). In conclusion, enhancers seem to fall into classes, which behave in similar ways with particular promoters, and the molecular compatibility that appears to tune PolII initiation rates seems to be mediated by the promoter motifs present in an enhancer-specific manner.

## Discussion

We dissected the *kni* gene locus as a case study of the role of multiple promoters in controlling a single gene’s transcription dynamics. Synthetic enhancer-promoter reporters allowed us to measure the ability of *kni* enhancer-promoter pairs to drive expression in the absence of complicating factors like promoter or enhancer competition. Using these reporters, we found that some promoters are broadly compatible with many enhancers, whereas others only drive high levels of expression with some enhancers. A detailed analysis of the transcription dynamics of these reporters indicates that the molecular compatibility of the proteins recruited to the enhancer and promoter tune expression levels by altering the initiation rate of transcriptional bursts.

In the context of the whole locus, we found that some enhancer-promoter pairs drive lower expression than their corresponding synthetic reporters, due to the effects of promoter and enhancer competition, distance, or other factors. In fact, while the synthetic reporters indicate that both promoters can drive similarly high levels of expression in the anterior, in the locus, promoter 1 drives most of the expression, with promoter 2 supporting some low levels of expression in the absence of promoter 1. In the posterior, both promoters appear to be necessary to achieve wildtype levels of expression with enhancer competition leading to sub-additive expression. By mutating promoter motifs in the synthetic enhancer-reporter constructs, we found that the effects of promoter motif mutations fall into two different classes, depending on the enhancer that is paired with the promoter. This suggests that there may be several discrete ways that a promoter can be activated by an enhancer, depending on the proteins recruited to each. Returning to our original hypotheses to explain the presence of two promoters in a single locus, we find that both differing enhancer-promoter preferences and a need for expression robustness in the face of promoter mutation may play a role.

Our work has highlighted the importance of both of *kni’s* promoters. Previous studies have almost exclusively focused on *kni*’s promoter 1 (Pankratz et al., 1992; Pelegri & Lehmann, 1994), which unexpectedly looks like a typical housekeeping gene promoter, with a dispersed shape and series of weak Inr sites (Vo Ngoc et al., 2017). It is *kni*’s promoter 2, with its focused site of initiation and composition of TATA Box, Inr, and DPE motifs, that looks like a canonical developmental promoter (Vo Ngoc et al., 2017). Interestingly, despite only discussing promoter 1, in practice, studies interrogating the behavior of multiple *kni* enhancers often included both promoters, as promoter 2 is found in a *kni* intron (Bothma et al., 2015; El-Sherif & Levine, 2016). Our analysis clearly demonstrates both promoters’ vital role in normal *kni* expression.

With these observations in mind, we wanted to determine the prevalence of a two-promoter structure, with one broad and one sharp. To do so, we used the RAMPAGE data set, which includes a genome-wide survey of promoter usage during the 24 hours of *Drosophila* embryonic development (P. J. Batut & Gingeras, 2017) and cross-referenced these promoters with those in the Eukaryotic Promoter Database, which is a collection of experimentally validated promoters (Dreos, Ambrosini, Groux, Cavin Périer, & Bucher, 2016). We found that 13% of embryonically expressed genes have at least two promoters. When we considered the two most commonly used promoters, there is a clear trend of a broader primary (most used) promoter (median = 91bp) and a sharper secondary promoter (median = 42bp) (Figure S1C). This trend is still present if the genes are split into developmental and housekeeping genes, with developmental promoters (median = 43bp) generally more focused than housekeeping promoters (median = 90bp), as expected (Figure S1D and E). Among the primary promoters of developmental genes, 58% consist of a series of weak Inr sites, much like *kni* promoter 1. This suggests that this promoter shape and motif content in developmental promoters may be more common than previously expected and should be explored.

There is growing evidence that promoter motifs play a role in modulating different aspects of transcription dynamics. However, the role of each motif can vary from one locus to the next. In the “TATA-only” *Drosophila snail* promoter, the TATA Box affects burst size by tuning burst duration (Pimmett et al., 2021). In the mouse PD1 proximal promoter, which consists of a CAAT Box, TATA Box, Sp1, and Inr motif, the TATA box may tune burst size and frequency (Hendy, Campbell, Weissman, Larson, & Singer, 2017). A study of a synthetic *Drosophila* core promoter and the *ftz* promoter found that the TATA box tunes burst size by modulating burst amplitude and that Inr, MTE, and DPE tune burst frequency (Yokoshi et al., 2021). TATA Box also appears to be associated with increased expression noise, as TATA-containing promoters tend to drive larger, but less frequent transcriptional bursts (Ramalingam et al., 2021). In contrast to TATA Box, Inr appears to be associated with promoter pausing, e.g. by adding a paused promoter state in the Inr-containing *Drosophila Kr* and *Ilp4* promoters (Pimmett et al., 2021). In fact, a Pol II ChIP-seq study indicates that paused developmental genes appear to be enriched for GAGA, Inr, DPE, and PB motifs (Ramalingam et al., 2021).

Similarly, the TFs bound at enhancers can affect transcription dynamics in diverse ways. Exploration of the role of TFs in modulating burst properties has indicated that BMP and Notch can tune burst frequency and duration, respectively (Falo-Sanjuan, Lammers, Garcia, & Bray, 2019; Hoppe et al., 2020; Lee, Shin, & Kimble, 2019). Work that considers both the promoters and enhancer simultaneously have come to differing conclusions. Work in human Jurkat cells, wherein 8000 genomic loci were integrated with one of three promoters, showed that burst frequency is modulated at weakly expressed loci and burst size modulated at strongly expressed loci (Dar et al., 2012). Work in *Drosophila* embryos and in mouse fibroblasts and stem cells suggest that stronger enhancers produce more bursts, and promoters tune burst size (Fukaya, Lim, & Levine, 2016; Larsson et al., 2019). On the whole, this work indicates that promoter motifs and the TFs binding enhancers can act to tune burst properties in a myriad of ways. Given the wide range of possibilities, it is likely that setting, i.e. the combination of promoter motifs and the interacting enhancers, is particularly important in determining the resulting transcription dynamics.

Our work supports this notion. Notably, eliminating the TATA Box and DPE from promoter 2 seems to reinforce the idea that we have two classes of enhancers that behave in distinct ways with these promoters due to the different TFs bound at these enhancers. We find that polymerase initiation rate is a key property tuned by the molecular compatibility of the proteins recruited to the enhancer and promoter. Our observation is in contrast to previous studies in which PolII initiation rate seems constant despite swapping two promoters with different motif content or altering BMP levels or the strength of TF’s activation domains (Hoppe et al., 2020; Senecal et al., 2014) and is tightly constrained for gap genes (Zoller, Little, & Gregor, 2018). We suggest that the differences we see in our work, where initiation rate depends on molecular compatibility, versus other work, where initiation rate is controlled by other factors, again reinforces the idea that the role of any particular promoter motif or TF binding site can be highly context dependent.

Together, ours and previous work demonstrate that deriving a general set of rules to predict transcription dynamics from sequence is a challenge because the space of promoter motif content and enhancer TF binding site arrangements is enormous. The proteins recruited to both promoters and enhancers can combine to make transcriptional complexes with different constituent proteins, post-translational modifications, and conformations, which may even vary as a function of time. Due to the vast possibility space and context-dependent rules, we have likely only scratched the surface of how promoter motifs or enhancers can modulate burst properties, suggesting a field rich for future investigation.

## Acknowledgements

The authors wish to thank Leonila Lagunes, Srikiran Chandrasekaran, and all the Wunderlich lab members for helpful comments on the manuscript and Ali Mortazavi, Kyoko Yokomori, Kevin Thornton, and Rahul Warrior for useful discussion on the project. The authors thank Flo Ramirez for data analysis that inspired some of this work.

## Funding

This work is supported by NIH-NICHD R00 HD073191 and NIH-NICHD R01 HD095246 (to ZW) and the US DoE P200A120207 and NIH-NIBIB T32 EB009418 (to LL).

## Conflicts of Interest

None to report.

## Materials and Methods

### Datasets used in this study

The experimentally validated promoters and their experimentally determined transcription start sites (TSSs) were obtained from the Eukaryotic Promoter Database (EPD) New (Dreos et al., 2016). They were cross-referenced with the RNA Annotation and Mapping of Promoters for Analysis of Gene Expression (RAMPAGE) data obtained from five species of *Drosophila* (P. J. Batut & Gingeras, 2017) to form a high-confidence set of promoters for which promoter usage during development could be evaluated. Single embryo RNA-seq obtained by Lott, et al. was indexed (with a *k* of 17 for an average mapping rate of 96%) and quantified using Salmon v0.12.01. The resulting transcript-specific data was used to further resolve *kni* promoter usage during nuclear cycle 14 (Lott et al., 2011; Patro, Duggal, Love, Irizarry, & Kingsford, 2017). Housekeeping genes were defined as in Corrales, et al. where genes were defined as housekeeping if their expression exceeded the 40^th^ percentile of expression in each of 30 time points and conditions using RNA-seq data collected by modEncode (Corrales et al., 2017) and a list of these can be found in the Supplementary Materials (File S1).

To study TF-promoter motif co-occurrence, we collected a total of ∼1000 enhancer-gene pairs expressed during development in *Drosophila*. The majority were identified by traditional enhancer trapping (REDfly & CRM Activity Database 2, or CAD2) and consist of non-redundant experimentally characterized enhancers (Bonn et al., 2012; Halfon, Gallo, & Bergman, 2008). About 15% were identified through functional characterization of ∼7000 enhancer candidates using high throughput *in situ* hybridization (Vienna Tile, or VT); these VT enhancers have been limited to those expressed during stages 4-6. The remaining 1% of enhancer-gene pairs have been identified through 4C-seq (Ghavi-Helm et al., 2014) and are active 3-4 hours after egg laying (stages 6-7). A list of these enhancer-promoter pairs and their coordinates can be found in the Supplementary Materials (File S2).

### Motif prediction in promoters and enhancers

For enhancers, TF binding site prediction was performed using Patser (Hertz & Stormo, 1999) with position weight matrices (PWMs) from the FlyFactor Survey (Zhu et al., 2011) and a GC content of 0.406. Each element in the PWM was adjusted with a pseudocount relative to the intergenic frequency of the corresponding base totaling 0.01. For TFs that had multiple PWMs available, PWMs built from the largest number of aligned sequences were chosen; that of Stat92E was taken from an older version of the FlyFactor Survey. For promoters, the transcription start clusters (TSCs) (P. J. Batut & Gingeras, 2017) and the adjoining ± 40bp were scanned for Inr, TATA Box, DPE, MTE, and TCT motifs using ElemeNT and the PWMs from (Sloutskin et al., 2015).

### Evaluation of total binding capacity of enhancers

Total binding capacity is a measure of the cumulative ability of an enhancer to bind a TF, and thus it takes into account the binding affinity of every *w*-mer in the enhancer for a TF binding site of length *w* (Wunderlich et al., 2012). To calculate the total binding capacity, we start by computationally scoring each possible site in the enhancer for the motifs of TFs regulating early axis specification. Taking the exponential of the score, normalizing this exponential by the enhancer length *l*, and summing these values gives us an overall binding capacity for each enhancer and TF combination, which is roughly equal to the sum of the probabilities that a TF is bound to each potential site in the enhancer.

Hence, we use the following formula

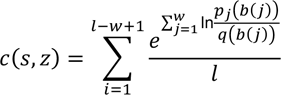

to calculate the total binding capacity *c* of a given sequence *s* for a given TF *z* (Wunderlich et al., 2012). Here, *l* is the length of the sequence being considered, *w* is the width of the PWM of the TF, *b*(*i*) is the base at position *i* of the sequence, *p*&(*b*) is the frequency of seeing base *b* at position *j* of the PWM, and *q*(*b*) is the background frequency of base *b*. Note that 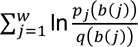 is equivalent to the score given to the *w*-mer at position *i* in the sequence calculated using Patser, as described above (Hertz & Stormo, 1999).

### Selection of enhancers to study

*knirps* enhancers expressed in the blastoderm were identified using REDfly (Halfon et al., 2008), and the shortest, non-overlapping subset of enhancers was obtained using SelectSmallestFeature.py available at the Halfon Lab GitHub (https://github.com/HalfonLab/UtilityPrograms). The enhancers in this subset were categorized by the expression patterns they drove, and a representative enhancer was picked from each of these categories.

### Generation of transgenic reporter fly lines

As described in Fukaya, et al., the four *kni* enhancers were each cloned into the pBphi vector, directly upstream of *kni* promoter 1, 2 or 2ΔTATAΔDPE; 24 MS2 repeats; and a *yellow* reporter gene (Fukaya et al., 2016). Similarly, the *kni* locus and its promoter knockouts (Δp1 and Δp2) were each cloned into the pBphi vector, directly upstream of 24 MS2 repeats and a *yellow* reporter gene by Applied Biological Materials (Richmond, BC, Canada). We defined kni_-5 as chr3L:20699503-20700905(–), kni_proximal_minimal as chr3L:20694587-20695245(–), kni_KD as chr3L:20696543-20697412(–), VT33935 as chr3L:20697271-20699384(–), promoter 1 as chr3L:20695324-20695479(–), promoter 2 as chr3L:20694506-20694631(–), and the *kni* locus as chr3L:20693955-20701078(–), using the *Drosophila melanogaster* dm6 release coordinates. Promoter motif knockouts (for p2ΔTATAΔDPE and locus Δp2) involved making the minimal number of mutations that would both inactivate the motif and introduce the fewest new motifs or TF binding sites (TATA: TA<u>T</u>AT<u>A</u>TATC > TA<u>G</u>AT<u>G</u>TATC, Inr: TC<u>A</u>GTT > TC<u>G</u>GTT, and DPE: A<u>G</u>A<u>T</u>CA > A<u>T</u>A<u>C</u>CA). The locus Δp1 construct involved replacing promoter 1 with a region of the lambda genome predicted to have the minimal number of relevant TF binding sites. The precise sequences for each reporter construct are given in a series of GenBank files included in the Supplementary Materials (File S3 – 18).

Using phiC31-mediated integration, each reporter construct was integrated into the same site on chr2L by injection into yw; PBac{y[+]-attP-3B}VK00002 (BDRC stock #9723) embryos by BestGene Inc (Chino Hills, CA). To visualize MS2 expression, female flies expressing RFP-tagged histones and GFP-tagged MCP (yw; His-RFP/Cyo; MCP-GFP/TM3.Sb) were crossed with males containing one of the MS2 reporter constructs.

### Sample preparation and image acquisition

As in Garcia et al., live embryos were collected prior to nuclear cycle 14 (nc14), dechorionated, mounted with glue on a permeable membrane, immersed in Halocarbon 27 oil, and put under a glass coverslip (Garcia et al., 2013). Individual embryos were then imaged on a Nikon A1R point scanning confocal microscope using a 60X/1.4 N.A. oil immersion objective and laser settings of 40uW for 488 nm and 35uW for 561 nm. To track transcription, 21 slice Z-stacks, at 0.5 um steps, were taken throughout nc14 at roughly 30s intervals. To identify the Z-stack’s position in the embryo, the whole embryo was imaged at the end of nc14 at 20x using the same laser power settings. To quantify expression along the AP axis, each transcriptional spot’s location was placed in 2.5% anterior-posterior (AP) bins across the length of the embryo, with the first bin at the anterior of the embryo. Embryos were imaged at ambient temperature, which was on average 26.5°C.

### Burst calling and calculation of transcription parameters

Tracking of nuclei and transcriptional puncta was done using a version of the image analysis MATLAB pipeline downloaded from the Garcia lab GitHub repository on January 8, 2020 and described in Garcia et al (Garcia et al., 2013). For every spot of transcription imaged, background fluorescence at each time point is estimated as the offset of fitting the 2D maximum projection of the Z-stack image centered around the transcriptional spot to a gaussian curve, using MATLAB *lsqnonlin*. This background estimate is subtracted from the raw spot fluorescence intensity. The resulting fluorescence traces across nc14 are then smoothed by the LOWESS method with a span of 10%. These smoothed traces are then used to quantify transcriptional properties and noise. Traces consisting of fewer than three timeframes are not included in the calculations.

To quantify the transcription properties of interest, we used the smoothed traces to determine at which time points the promoter was “on” or “off” (Waymack et al., 2020). A promoter was considered “on” if the slope of its trace, i.e. the change in fluorescence, between one point and the next was greater than or equal to the instantaneous fluorescence value calculated for one mRNA molecule (F_RNAP_, described below). Once called “on”, the promoter is considered active until the slope of the fluorescence trace becomes less than or equal to the negative instantaneous fluorescence value of one mRNA molecule, at which point it is considered inactive until the next time point it is called “on”. The instantaneous fluorescence of a single mRNA was chosen as the threshold because we reasoned that an increase in fluorescence greater than or equal to that of a single transcript is indicative of an actively producing promoter, just as a decrease in fluorescence greater than that associated with a single transcript indicates that transcripts are primarily dissociating from, not being newly initiated at, this locus. Visual inspection of fluorescence traces agreed well with the burst calling produced by this method (Figure S4) (Waymack et al., 2020).

Using these smoothed traces and “on” and “off” time points of promoters, we measured burst size, burst frequency, burst duration, polymerase initiation rate, and noise. Burst size is defined as the integrated area under the curve of each transcriptional burst, from one “on” frame to the next “on” frame, with the value of 0 set to the floor of the background-subtracted fluorescence trace (Figure S4C). Frequency is defined as the number of bursts in nc14 divided by time between the first time the promoter is called active and 50 min into nc14 or the movie ends, whichever is first (Figure S4E). The time of first activity was used for frequency calculations because the different enhancer constructs showed different characteristic times to first transcriptional burst during nc14. Duration is defined as the amount of time occurring between the frame a promoter is considered “on” and the frame it is next considered “off” (Figure S4F). Polymerase initiation rate is defined as the slope at the midpoint between the frame a promoter is considered “on” and the frame it is next considered “off” (Figure S4G). The temporal coefficient of variation of each transcriptional spot *i*, was calculated using the formula:

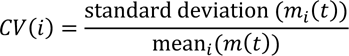

where *m_i_*(*t*) is the fluorescence of spot *i* at time *t*. For these, and all other measurements, we control for the embryo position of the fluorescence trace by first individually analyzing the trace and then using all the traces in each AP bin (anterior-posterior; the embryo is divided into 41 bins each containing 2.5% of the embryo’s length) to calculate summary statistics of the transcriptional dynamics and noise values at that AP position.

All original MATLAB code used for burst calling, noise measurements, and other image processing are available at the Wunderlich Lab GitHub (Waymack et al., 2020) with a copy archived at https://github.com/elifesciences-publications/KrShadowEnhancerCode. Updates to include calculations of polymerase initiation rate are also available at the Wunderlich Lab GitHub (https://github.com/WunderlichLab).

### Conversion of integrated fluorescence to mRNA molecules

To convert arbitrary fluorescence units into physiologically relevant units, we calibrated our fluorescence measurements in terms of mRNA molecules. As in Lammers et al., for our microscope, we determined a calibration factor, α, between our MS2 signal integrated over nc13, F_MS2_, and the number of mRNAs generated by a single allele from the same reporter construct in the same time interval, N_FISH_, using the *hunchback* P2 enhancer reporter construct (Garcia et al., 2013; Lammers et al., 2020). Using this conversion factor, we calculated the integrated fluorescence of a single mRNA (F_1_) as well as the instantaneous fluorescence of an mRNA molecule (F_RNAP_). For our microscope, F_RNAP_ is 379 AU/RNAP, and F_1_ is 1338 AU/RNAP·min. We can use this values to convert both integrated and instantaneous fluorescence into total mRNAs produced and number of nascent mRNAs present at a single time point, by dividing by F_1_ and F_RNAP_, respectively.

### Regression modeling and statistical analysis

To quantify the effect of enhancer, promoter, and interaction terms on burst parameters, we considered models of the form

*g*(*Y*) = enhancer + promoter + (enhancer × promoter)

where *Y* is the burst property of interest and *g* is the link function (Figure 4A). Model selection involved considering (1) the type of model, (2) the distribution that best fit the burst property data and (3) the appropriate predictors to include. We approached model selection with no specific expectations, opting to use generalized linear models (GLMs) because they were not much improved upon by adding random effects (GLMMs) and because they fit the data better than linear models (LMs).

Similarly, the appropriate distribution for each burst property was determined by fitting various distributions to the data and comparing their goodness-of-fit. As expected, total RNA produced and burst size (in transcripts per burst) were best described by a negative binomial distribution, as has been commonly used to describe count data. For the other burst properties, for which the appropriate distribution was less clear, we found that burst frequency was best fit by the Weibull distribution and burst duration and initiation rate were best fit by the gamma distribution. These choices were supported by the lower AIC values produced when comparing them to models using alternative distributions. They also seem reasonable given examples of other applications of these distributions. To keep the interpretation consistent across models, we chose to use an identity link function for all models (Figure 4B); using the canonical link functions associated with each of these distributions produced the same trends (Figure S5).

The predictors we included were the enhancer and promoter and any interaction terms between the enhancer and promoter. In each case, dropping the interaction terms produced higher AIC values, suggesting that the interaction terms are important and should not be dropped by the model.

To determine any significant differences in mean expression levels, we performed Welch’s *t*-tests, and to determine if any predictors led to significant differences in burst duration, we performed a multivariate ANOVA. To quantify the variability explained by different predictors, we calculated the Cragg and Uhler pseudo R-squared measures of the model including only the predictor in question and divided by that of the full model described above.

### Data Availability Statement

Transgenic fly strains and plasmids are available upon request. Supplementary File S1 contains the gene names, the dm6 release coordinates, and the FlyBase numbers (FBgns) that matched to the gene names and coordinates (Corrales et al., 2017). File S2 contains DNA sequences of the enhancers and promoters used in the computational analysis presented in Figure S2. Files S3-18 contain GenBank files describing the plasmids used to make all the transgenic fly strains produced for this work.

---

**Figure S1.**
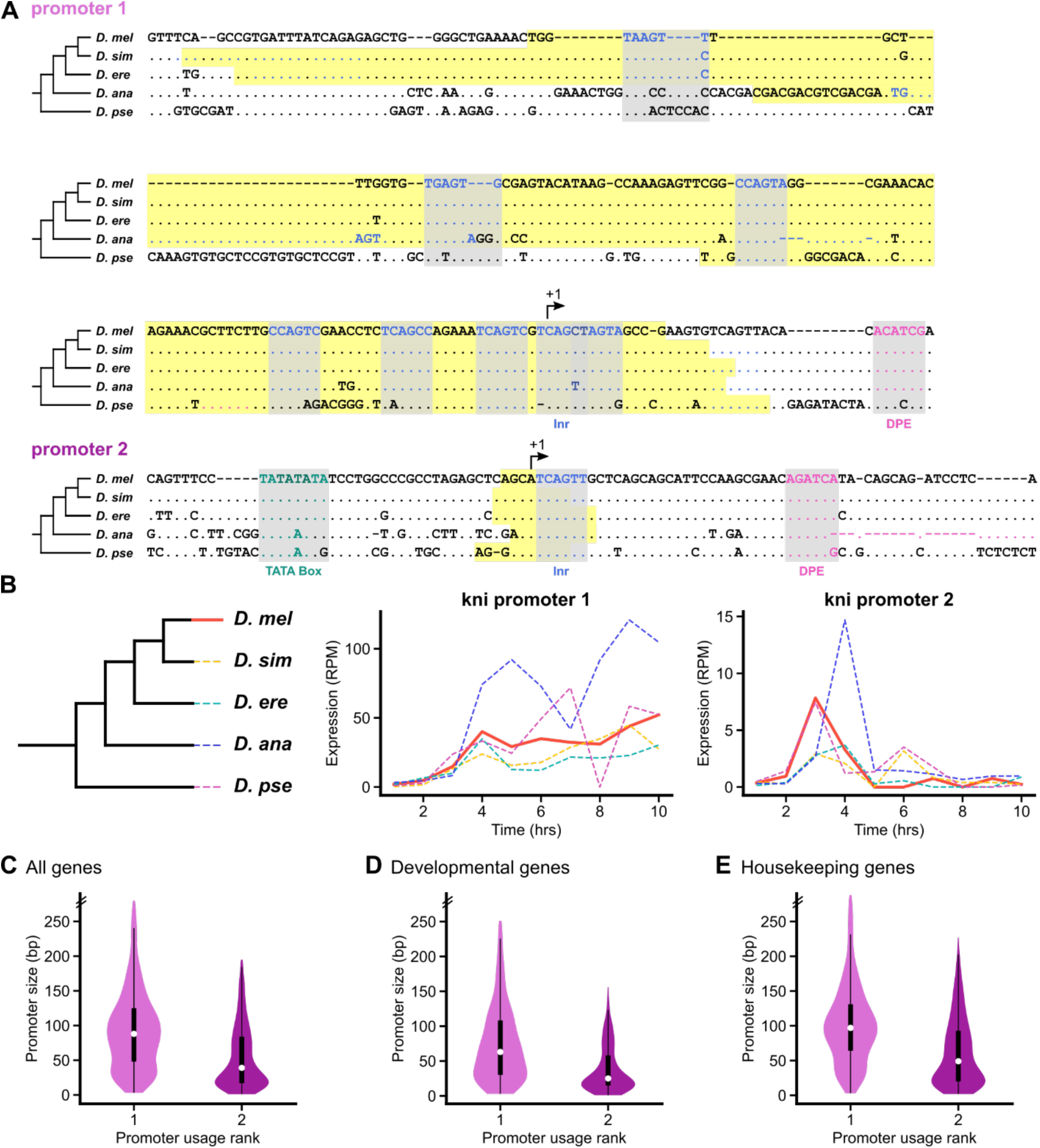
The *knirps* promoters show sequence and functional conservation, and this two-promoter structure is prevalent among genes expressed during development. (**A**) Both *kni* promoters are aligned with the orthologous sequences in four other *Drosophila* species, with dashes (-) representing unaligned sequence and dots (.) indicating matching base pairs. There is remarkable sequence conservation, with the core promoter motifs preserved across all five species. The highlighted regions represent transcription start clusters (TSCs), identified by Batut, et al (P. J. Batut & Gingeras, 2017) as regions of statistically significant clustering of cDNA 5’ ends. (**B**) *Kni* promoter activity over the first 10 hours of development is reasonably consistent across five species of *Drosophila*, with promoter 1 generally being used more than promoter 2. Specifically, note that both promoters are used in nuclear cycle 14 (2-3 hours) in all five species. **(C – E)** For developmentally expressed genes with multiple promoters that are represented in both the Eukaryotic Promoter Database and the Batut *et al*. RAMPAGE data (P. J. Batut & Gingeras, 2017; Dreos, Ambrosini, Cavin Périer, & Bucher, 2013), violin plots of the two most used promoters, with the primary promoter (most used) in light purple and the secondary promoter (second most used) in dark purple. The black boxes span the lower to upper quartiles, with the white dot within the box indicating the median. Whiskers extend to 1.5*IQR (interquartile range) ± the upper and lower quartile, respectively. The double hash marks on the axes indicate that 95% of the data is being shown. **(C)** When the two most used promoters of genes expressed in embryogenesis (*n* = 1177) are plotted, the size of primary promoters is significantly larger than that of the secondary promoter. **(D)** When limited to promoters of developmentally controlled genes – genes whose expression pattern varies considerably as a function of developmental time -- (*n* = 387) this trend of larger primary promoters is maintained, though on average, these promoters are sharper that those of the whole gene set in panel C. **(E)** When limited to promoters of housekeeping genes (*n* = 790), this trend of larger primary than secondary promoters is also maintained, though on average, these promoters are still broader than those of developmentally controlled genes.

**Figure S2.**
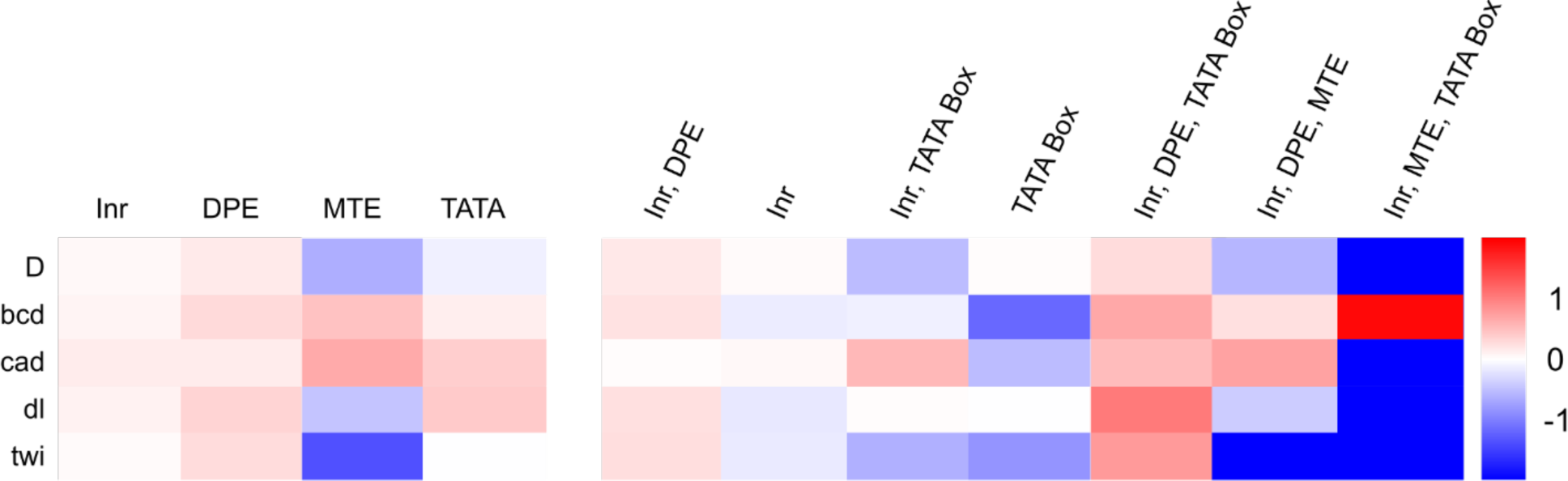
TFs show preferences for certain core promoter motifs. To identify patterns of TF-core promoter motif co-occurrence, we calculated the fold enrichment of core promoter elements associated with TF-target genes. The left heatmap shows the log fold-enrichment over background of the frequency of the core promoter motif (columns) for the set of promoters associated with enhancers controlled by the TF (rows). The right heatmap shows the log fold-enrichment over background of the frequency of the motif combination (columns) for the set of promoters associated with enhancers controlled by the TF (rows). For example, this means that column 1 (Inr) in the left heatmap shows enrichment of any promoters that contain Inr regardless of any other promoter motifs they might contain, whereas column 2 (Inr) in the right heatmap shows enrichment of promoters with only Inr and no other core promoter motifs.

**Figure S3.**
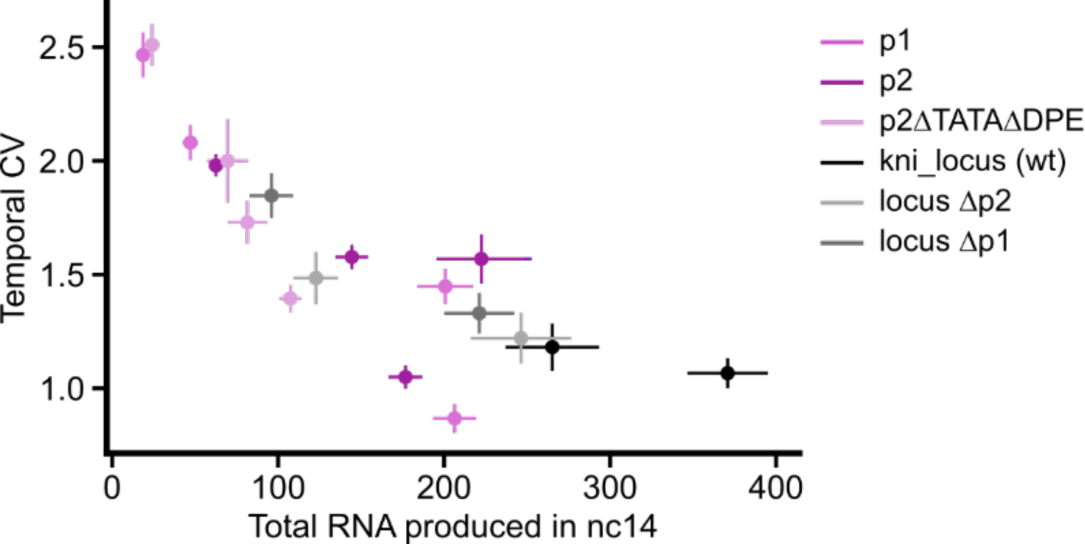
Noise is inversely correlated with total RNA produced. To examine the relationship between temporal coefficient of variation (CV) and activity of each construct, we plotted the mean temporal CV against the total RNA produced in nc14 at the anterior-posterior bin of maximum expression (22% and 63%) for the anterior band and the posterior stripe, respectively, with the error bars representing 95% confidence intervals. There is a clear trend of CV decreasing with increased total RNA produced though there are examples where constructs with the same promoter can produce higher noise than others with similar output levels, suggesting that promoters do not solely dictate noise levels.

**Figure S4.**
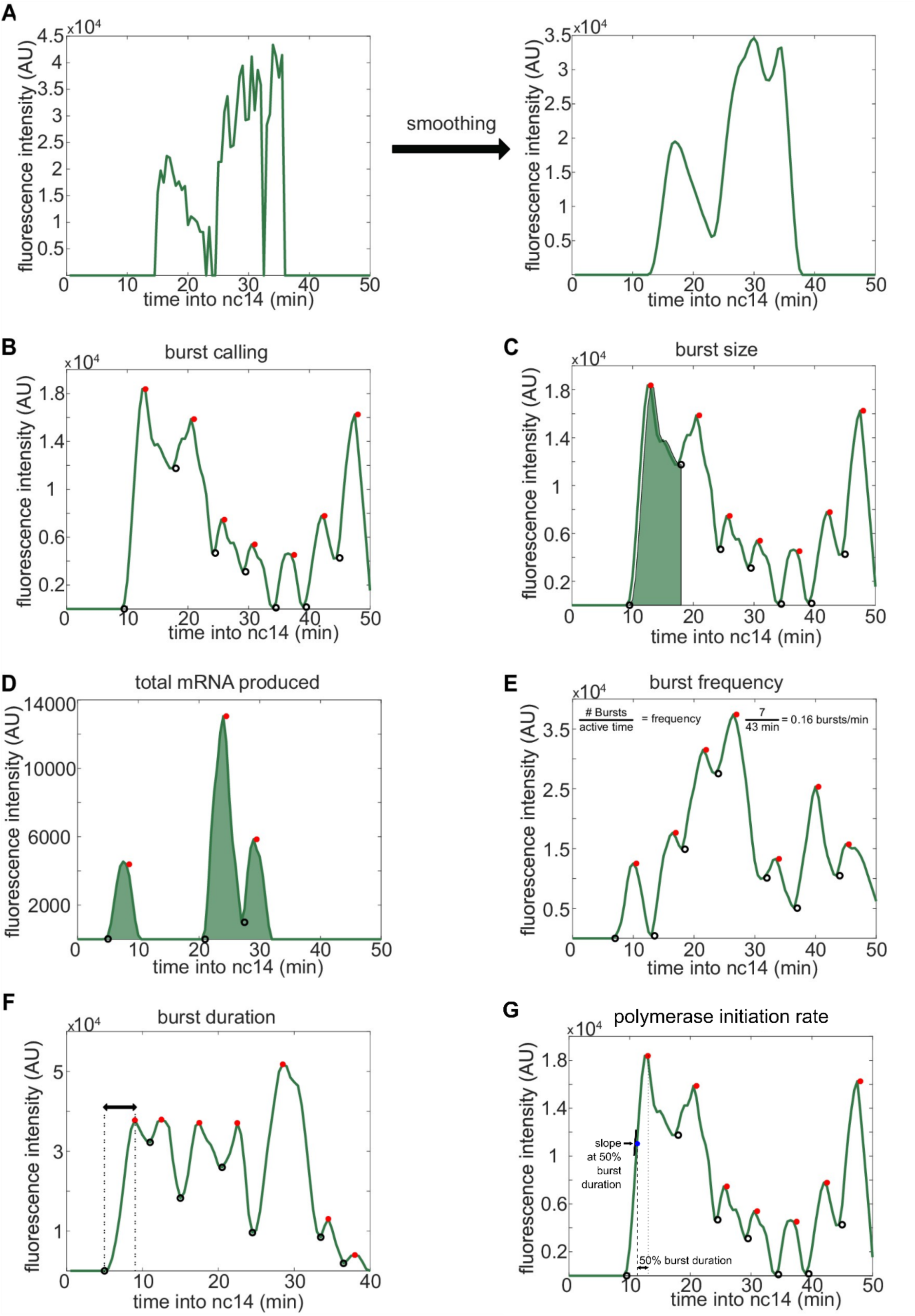
Visual inspection of burst calling algorithm. This figure is adapted from Waymack, et al. with one additional panel **(G)** added (Waymack et al., 2020). To quantify the burst properties of interest (burst size, burst frequency, burst duration, and polymerase initiation rate), we began by smoothing individual fluorescence traces using the LOWESS method with a span of 10%. Periods of promoter activity or inactivity were then determined based on the slope of the fluorescence trace. **(A)** Example of smoothing transcriptional traces. **(B)** Fluorescence trace of a single punctum during nc14. Open black circles indicate time points where the promoter has turned “on”, filled red circles indicate time points where the promoter is identified as turning “off”. **(C)** Transcriptional trace with the green shaded region under the curve used to calculate the size of the first burst. This area of this region is calculated using the trapz function in MATLAB and extends from the time point the promoter is called “on” until the next time it is called “on”. Panels **(D – F)** show additional representative fluorescence traces of single transcriptional puncta during nc14. **(D)** A trace with the entire region under the curved shaded green represents the area used to calculate the total amount of mRNA produced. This area is calculated using the trapz function in MATLAB extends from the time the promoter is first called “on” until 50 min into nc14 or the movie ends, whichever comes first. **(E)** Burst frequency is calculated by dividing the number of bursts that occur during nc14 by the length of time from the first time the promoter is called “on” until 50 min into nc14 or the movie ends, whichever comes first. **(F)** Burst duration is calculated by taking the amount of time between when the promoter is called “on” and it is next called “off”. **(G)** Polymerase initiation rate is calculated by taking the slope of the smoothed fluorescence race at the midpoint between when the promoter is called “on” and it is next called “off”.

**Figure S5.**
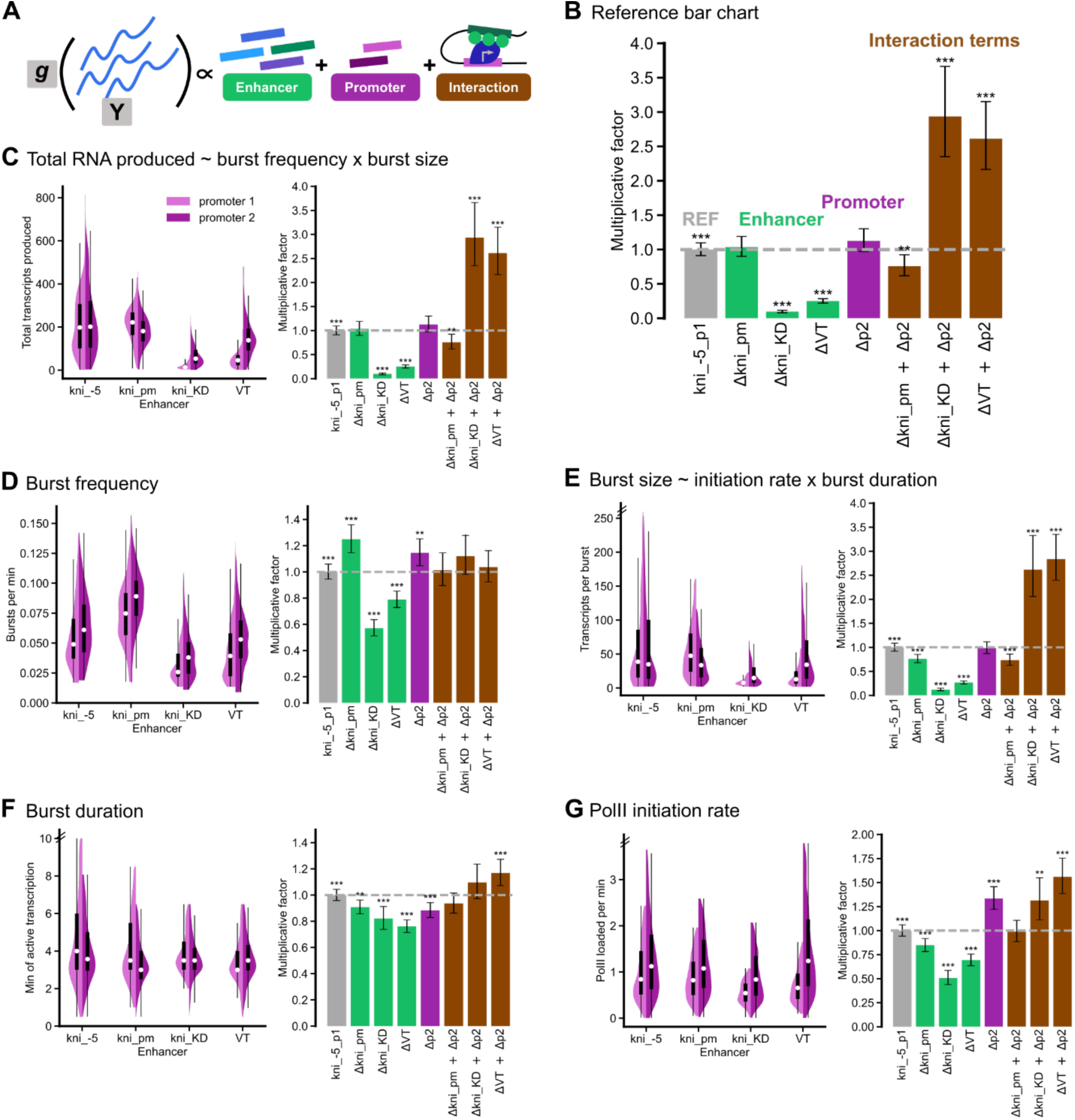
Using canonical link functions gives the same results. Here, we show the results from the generalized linear models (GLMs) when using the log link function instead of the identity link function, which was used in Figure 4. **(A)** To parse the effects of the enhancer, the promoter, and their interactions on all burst properties, we built GLMs. Y represents the burst property under study, g is the link function, and the enhancers, promoters, and their interaction terms are the explanatory variables. The coefficients of each of these explanatory variables is representative of that variable’s contribution to the total value of the burst property. **(B)** All burst property data was taken from the anterior-posterior bin of maximum expression (22% and 63%) for the anterior band and the posterior stripe, respectively. We exponentiate the coefficients and the 95% confidence intervals for each independent variable to invert the log link function and call these quantities the “multiplicative factors.” Performing this conversion yields a multiplicative relationship between our response variable (the burst property) and our explanatory variables. The reference construct (kni_-5-p1) has been set to 1 such that multiplying the relevant multiplicative factors gives you the value that, if multiplied by the reference construct value, will gives you the average value of the burst property for a particular construct. These factors are plotted as a bar graph; * *p* < 0.05, ** *p* < 0.01, *** *p* < 0.001. The reference construct is represented in gray, and the effects of enhancer, promoter, and their interactions are represented in green, purple, and brown, respectively. Thus, the average value of the burst property for a particular construct, e.g. VT-p2, relative to the reference construct would be 0.73, which is the product of ΔVT = 0.25, Δp2 = 1.1, and ΔVT + Δp2 = 2.6. The average value of the burst property for VT-p2 would then be 0.73 × 205 = 150. Note that for simplicity, kni_proximal_minimal and VT33935 has been shortened to kni_pm and VT, respectively, in the following graphs. In panels **(C – G)**, (left) split violin plots (and their associated box plots) of burst properties for all eight constructs will be plotted with promoter 1 in light purple and promoter 2 in purple. The black boxes span the lower to upper quartiles, with the white dot within the box indicating the median. Whiskers extend to 1.5*IQR (interquartile range) ± the upper and lower quartile, respectively. (right) Bar graphs representing the relative contributions of enhancer, promoter, and their interactions to each burst property are plotted as described in **(B)**. The double hash marks on the axes indicate that 90% of the data is being shown. **(C)** Expression levels are mainly determined by the enhancer and the interaction terms. Some enhancers (kni_-5 and kni_proximal_minimal) appear to work well with both promoters; whereas, kni_KD and VT, which are bound by similar TFs, show much higher expression with promoter 2. **(D)** Burst frequency is dominated by the enhancer and promoter terms, with promoter 2 consistently producing higher burst frequencies regardless of enhancer. **(E)** Burst size, which is determined by both initiation rate and burst duration, is dominated by the enhancer and interaction terms. As **(F)** burst duration is reasonably consistent regardless of enhancer or promoter, differences in burst size are mainly dependent on differences in **(G)** PolII initiation rate, with this burst property as the main molecular knob affected by molecular compatibility.

